# Astroglial biophysics probed with a realistic cell model

**DOI:** 10.1101/336974

**Authors:** Leonid P. Savtchenko, Lucie Bard, Thomas P. Jensen, James P. Reynolds, Igor Kraev, Mikola Medvedev, Michael G. Stewart, Christian Henneberger, Dmitri A. Rusakov

**Affiliations:** UCL Institute of Neurology, University College London, UK; The Open University, Milton Keynes, UK; Institute of Cellular Neurosciences, University of Bonn, Germany; German Center of Neurodegenerative Diseases (DZNE), Bonn, Germany

## Abstract

Electrically non-excitable astroglia take up neurotransmitters, buffer extracellular K^+^ and generate Ca^2+^ signals that release molecular regulators of neural circuitry. The underlying machinery remains enigmatic, mainly because the nanoscopic, sponge-like astrocyte morphology has been difficult to access experimentally or explore theoretically. Here, we have systematically evaluated the multi-scale morphology of protoplasmic astroglia to construct a realistic multi-compartmental cell model that can be biophysically interrogated in NEURON computational environment. This approach has been implemented as an astrocyte-model builder ASTRO. As a proof of concept, we explored a hippocampal astrocyte reconstructed *in silico* against a battery of physiological and imaging experiments. This exploration has unveiled some basic features of astroglial physiology inaccessible empirically, such as the characteristic length of membrane voltage propagation, membrane effects of local glutamate transport, spatiotemporal dynamics of intracellular K^+^ redistribution, key Ca^2+^ buffering properties, and some basic relationships between free Ca^2+^ dynamics and experimental readout of fluorescent Ca^2+^ indicators.

---

Astroglia have emerged as an essential contributor to neural circuit signalling in the brain. In addition to the well-established mechanisms of neurotransmitter uptake and extracellular K^+^ buffering, electrically passive astrocytes appear competent in transducing, integrating and propagating physiological signals using intracellular Ca^2+^ transients ^1–3^. Astroglial Ca^2+^ signals range from local hotspots to global Ca^2+^ waves, displaying a variety of dynamic ranges and time scales (reviewed in ^4, 5^). Tri-dimensional (3D) reconstructions of astroglia using electron microscopy (EM) have long revealed a fine system of nanoscopic processes ^6–10^ that pervade the entire cell expanse ^11–13^. Deciphering the cellular cascades and the biophysical mechanisms that shape Ca^2+^-dependent signalling in this sponge-like system has been a challenge ^14, 15^. A similar challenge pertains to our understanding of how the complex geometry of astrocytes controls intracellular ionic homeostasis during rapid local influx of K^+^, other ions, or water molecules ^16–23^.

In contrast, cellular mechanisms underpinning neuronal physiology have been explored and understood in great detail. This owes to the fact that rapid advances in patch-clamp electrophysiology and cell imaging in neurons have been recapitulated and mechanistically interpreted using realistic biophysical cell models *in silico*. The multi-compartmental modelling of various neuronal types developed in the NEURON computational environment^24, 25^ has been highly instrumental in advancing our knowledge about neural signal integration in the brain. There have also been numerous attempts to simulate astroglial function, based predominantly on a reductionist approach (recently reviewed in ^26, 27^). Aimed to answer a specific scientific question, such models would normally focus on kinetic reactions inside astroglia^28–30^, between astroglial and neuronal compartments ^16, 31–33^ or, alternatively, on the regulatory influences of astroglial signalling in neuronal networks ^34–36^. These studies have provided some important insights into the biophysical basis of astroglial activity and its adaptive roles. However, their scope would normally exclude complex 3D morphology of astrocytes, intracellular concentration gradients and heterogeneous molecular fluxes, uneven landscapes of cell membrane potential, or complex relationships between Ca^2+^ kinetics and Ca^2+^-sensitive fluorescence recordings. Thus, to date there have been no attempts to integrate cellular functions of astrocyte on multiple levels, in one entity *in silico*, with the realistic morphology playing a major role in shaping astroglial identity ^14^.

Our aim was therefore three-fold. First, to generate a modelling approach that recapitulates fine astroglial morphology on multiple scales while retaining full capabilities of biophysical simulations enabled by NEURON. We have therefore developed (MATLAB- and NEURON-based) computational algorithms and software that, firstly, use experimental data to recreate the space-filling architecture of nanoscopic astroglial processes, and, secondly, make this cell architecture NEURON-compatible. Our case study focused on the common type of hippocampal protoplasmic astroglia, which has been the main subject of studies focusing on synaptic plasticity and neuron-glia interactions ^37–40^. We have combined patch-clamp electrophysiology, two-photon excitation (2PE) imaging and 2PE spot-uncaging, fluorescence recovery from photobleaching (FRAP), viral transduction of astroglia-targeted Ca^2+^ indicators *in vivo*, and quantitative correlational 3D EM to systematically document the multi-scale morphology and some key physiological traits of these cells. Based on these experimental constrains, we have built a multi-compartmental 3D cell model that recapitulates known features of hippocampal astrocytes and is fully integrated into the NEURON environment. Correspondingly, we have equipped the latter with additional functionalities relevant to astroglial studies, such as: generation of nanoscopic processes that fill in tissue volume; options for incorporating endfoot and gap junctions; options to arrange extracellular glutamate spot-uncaging and localised extracellular K^+^ rises; menu for running 3D FRAP experiments; intracellular Ca^2+^ signal generation options; settings to replicate conditions of fluorescence imaging.

Our second objective was to implement this modelling approach as a user-friendly, interactive simulation instrument - cell model builder - capable of recreating and probing various types of astroglia *in silico*. Thus, we have integrated our algorithms and software as a modelling tool ASTRO, which enables an investigator to generate morphological and functional astroglial features at various scales (current version at https://github.com/LeonidSavtchenko/Astro).

Finally, as a proof of concept, we explored and probed the test-case model against experimental observations, aiming to understand some important aspects of astroglial physiology that cannot be accessed through direct experimental probing. In our case study focusing on *stratum radiatum* astroglia, we have therefore evaluated key electrogenic features of the cell membrane, basic aspects of intracellular K^+^ dynamics, the range of intracellular Ca^2+^ buffering capacity, and how the classical molecular machinery of Ca^2+^ ‘puffs’ and ‘sparks’ could explain the common observations in astroglia using fluorescence imaging of Ca^2+^-sensitive indicators. The results suggest that ASTRO could provide an important means for testing physiological hypotheses and causal interpretation of experimental observations pertinent to astrocytes.

## RESULTS

### Reconstructing gross morphology of live astroglia: stem tree

The gross morphology of CA1 astrocytes has long been documented ^11, 41, 42^ pointing to the overall cell radius of 30-50 μm, the somatic diameter of 7-15 μm, 4-9 primary processes ^11, 43^, and a main-branch ramification index (RI, probability of bifurcation per branch) of less than one ^44^. To elucidate the structural detail of CA1 astrocytes *in situ* we employed acute hippocampal slices. Individual astroglia were held in whole-cell, loaded with the bright morphological tracer Alexa Fluor 594 (Methods), and imaged using two-photon excitation (2PE; Fig. 1a,b), a procedure shown quantitatively to faithfully reveal fine cell morphology ^42^.

**Figure 1.**
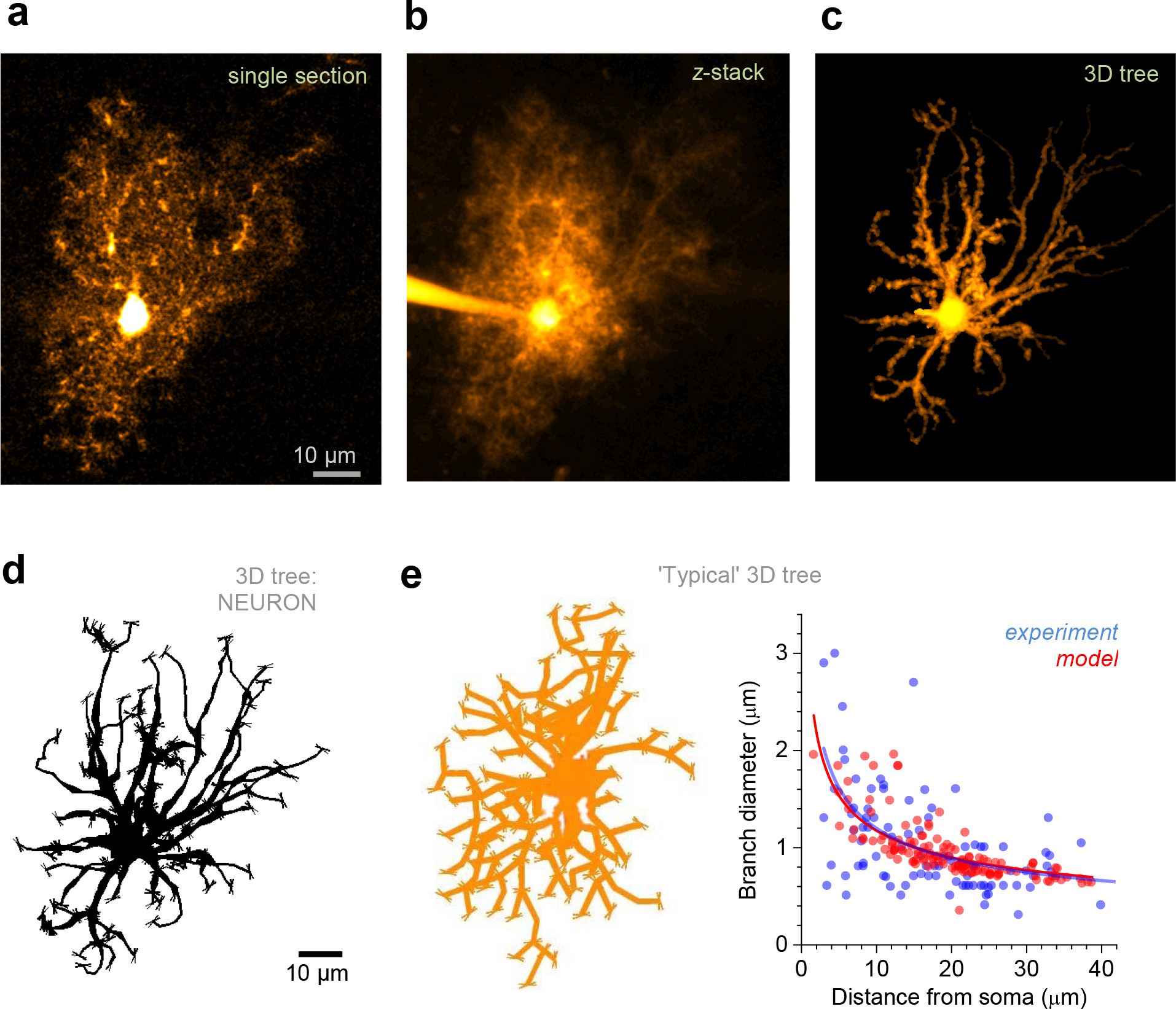
Reconstructing astroglial stem tree *in silico*. **a**, A characteristic image of CA1 astroglia, whole-cell load with Alexa Fluor 594 (λ_x_^2p^ = 800 nm), single optical section (*stratum radiatum*, depth of ~100 μm). **b**, Cell as in panel **a** shown as a full *z*-stack projection. **c**, Stem tree of astroglia shown in **a-b**, separated and reconstructed in 3D using NeuroTrace (Fiji ImageJ, NIH); 2D view of a 3D image. **d**, Astrocyte stem tree shown in panel c quantified, loaded and displayed in NEURON format using Vaa3D (Allen Institute); thin ‘buds’ indicate initial seeds for ‘planting, nanoscopic protrusions at a certain longitudinal density; 2D view. **e**,Diagram, ‘typical’ astrocyte stem tree built by modifying a library neurogliaform cell (2D view); plot, matching the branch diameters in the model (red) and in recorded astroglia (blue; n = 13 cells including 98 dendrites); solid lines, the best-fit dependence (power low, *y* = *a* ·*x*^*b*^) for the corresponding data scatters.

Our modelling strategy was to start with the principal branch structure (cell ‘stem tree’) which could be resolved in 2PE optical images (i.e., with a diameter above the diffraction limit, 0.3-0.5 μm; Fig. 1a,b) and 3D-reconstructed using computer tools that have been previously validated in neuronal studies. Here, we have employed two complementary approaches. In the first approach (Methods), the cell was imaged in a *z*-stack (Fig. 1a,b), individual images were corrected for the depth-dependent fluorescence signal decrease, identifiable stem-tree branches were traced semi-automatically and 3D-reconstructed using Simple Neurite Tracer (Fiji-ImageJ, NIH; Fig. 1c; Methods). This routine was similar to that of reconstructing 3D geometry of neuronal processes, as described widely in the literature. To store the recorded structure data in a NEURON-compatible format (Fig. 1d) we used Vaa3D (Allen Institute). Alternatively, the entire 3D reconstruction procedure could be carried out using commercially available Neurolucida (MBI), which provides a NEURON-compatible output.

The second approach was to build the stem tree of the ‘typical’ (or ‘average’) astrocyte from the population under study. First, the stem tree structure was taken from a digital library: in our case study we used a library neurogliaform cell (P32-DEV136, database http://neuromorpho.org), which displays gross morphology similar to that of the astrocyte (Fig. 1e, diagram). Next, the numbers and diameters of tree branches were adjusted to match experimental measurements from an experimental sample (in our case 13 astrocytes, 98 individual branches; Fig. 1e). The stem tree options also include settings to construct and explore an endfoot structure (Supplementary ASTRO User Guide, pp 20-21).

### The endfoot

The endfoot surrounding blood vessels is a key morphological and functional feature of most astrocytes ^45, 46^. Because its morphology varies enormously among cells, it would seem appropriate to use direct experimental 3D reconstructions (as in Fig. 1a-d) to incorporate this feature into the cell architecture. ASTRO provides a separate menu for constructing the endfoot, connecting it to the ‘main’ arbour, and populating it with the nanoscopic processes (Supplementary ASTRO User Guide, pp 20-21). All nanoscopic process structures and biophysical mechanisms available in the present model builder (see sections below) could be incorporated into the endfoot. However, as it represents a self-contained, highly specialised cell compartment, it will require a separate study to fully develop its realistic biophysical machinery and incorporate it in accord with experimental observations. Simulation tests in the present study focus therefore on the ‘main’ astrocyte arbour, which is thought to play a key role in astroglia-neuron interactions ^4, 5, 14, 47^. Nonetheless, parts of the main arbour may include processes that surround small blood vessels.

### Nanoscopic astroglial processes: experimental measurements

Once the cell stem tree was ready and NEURON-formatted, it was imported by ASTRO for the next step of morphological reconstruction: stochastic generation of nanoscopic processes. Whilst the stem tree branches can be traced in the light microscope, the bulk of astroglial morphology comprises irregularly shaped, nanoscopic processes that occur throughout the cell expanse and appear as a blur in optical images (Fig. 1a,b). 3D EM reconstructions have been used to detail the spatial organisation of these processes ^10, 12, 13, 42^. To systematically quantify such structures, we employed a correlational-EM approach based on our earlier 3D EM studies of astroglia ^10, 48^. Individual astrocytes were patched whole-cell in acute brain slices and, after cell filling with biocytine and subsequent DAB conversion, traced and reconstructed in a comprehensive serial-section survey (Methods; Supplementary Fig. 1). A recent EM study has suggested that chemical fixation via heart perfusion might lead to 30-35% volumetric tissue shrinkage and concomitant changes in fine astroglial morphology ^49^. To minimise such effects, we used rapid fixation of thin acute slices by submersion, which was earlier shown to cause much smaller hippocampal tissue shrinkage (linear shrinkage ~5%) ^50^ preserving the extracellular space volume fraction of ~0.12 (reviewed in ^51^) which was close to ~0.15 seen under rapid cryofixation ^49^.

Obtaining 3D reconstruction of the entire astrocyte at nanoscopic resolution is technically difficult and it may not necessarily represent the ‘typical’ cell. We therefore focused on the characteristic morphological features of 3D-reconstructed fragments sampled from multiple individual astrocytes from the CA1 population, as detailed earlier ^10^ (Fig. 2a). The aim of this exploration was to extract key morphometric characteristics of the nanoscopic astrocyte processes in order to replicate such ‘typical’ processes in the cell model.

**Figure 2.**
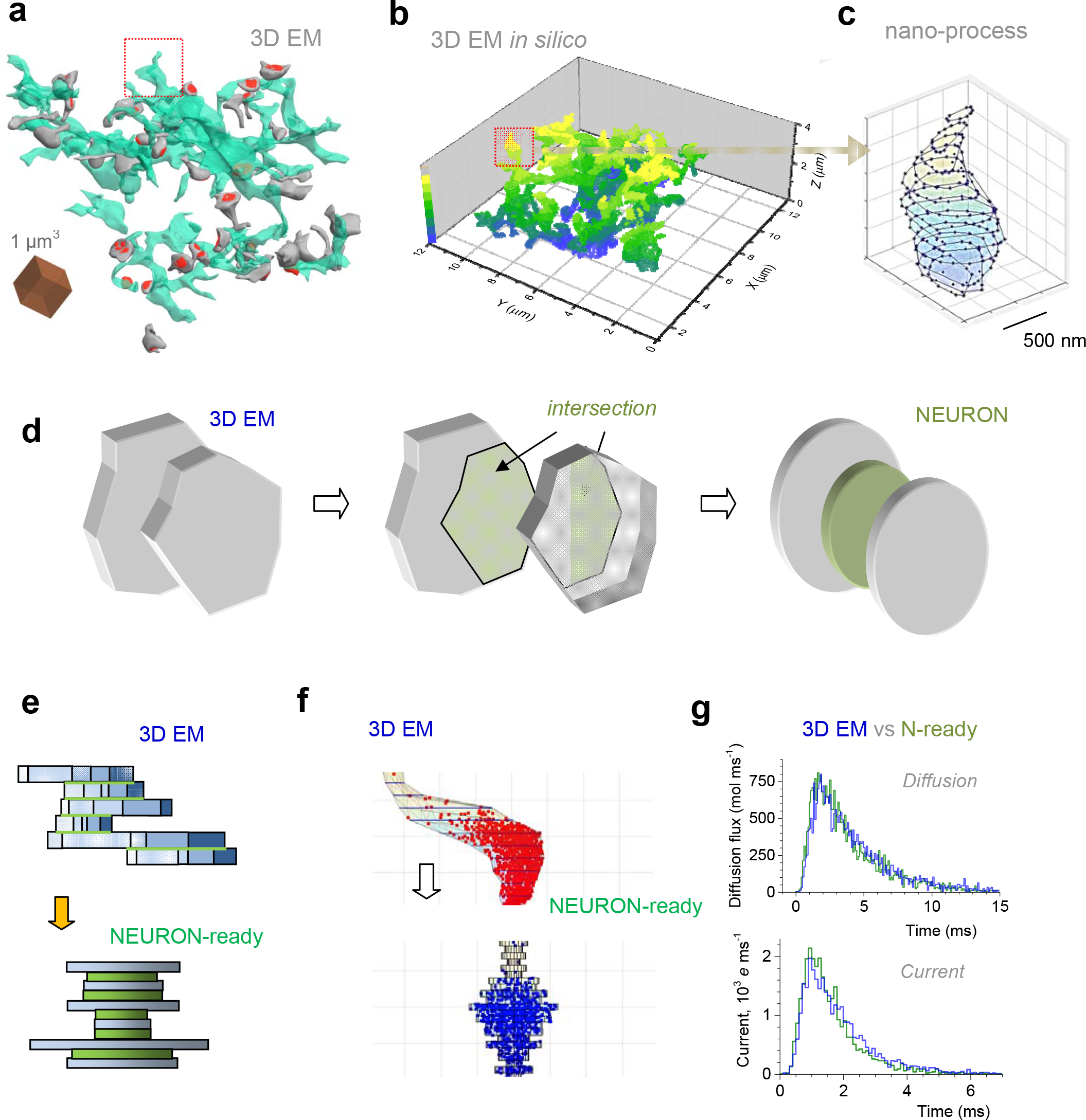
Documenting and reconstructing *in silico* nanoscopic astroglial protrusions. **a**, 3D EM reconstruction of an astrocytic fragment (green) and adjacent dendritic spines (grey) with postsynaptic densities (red), from serial sections in area CA1; surface rendering applied for clarity, as in ^10^; dotted square, fragment of interest. **b**, Fragment shown in a but with the surface point scatter delimiting the shape; false colour scale, z-depth as indicated; dotted square, fragment of interest (nanoscopic astroglial process) as in **a**. **c**, Astroglial process highlighted in **a-b** shown as a stack of serial polygonal sections (60 nm thick, to follow EM serial sections) delimited by the surface points. **d**, Transformation of the adjacent EM sections (*left;* grey polygonal slabs with base areas *S*_*i*_ and *S*_*i*+*1*_) with intersection area *T*_*i*_ (*middle*; green polygons) into NEURON-compatible two main (grey, ‘leaves’) and one transitional (green, ‘stalk’) cylindrical slabs with the corresponding base areas (*right*). **e**, Transformation of the 3D EM reconstructed process fragment (top panel) into NEURON-compatible cylinder section stacks (*bottom* panel); side view. Individual sections (blue, *top*) were transformed into ‘main’ cylinders of the same base area (blue, *bottom*). Green segments (top) depict adjacent surfaces between sections, which are represented by green ‘transitional’ cylinders (*bottom*), as detailed in **d**. **f**, A characteristic example of a 3D-EM reconstructed astroglial process represented by serial polygonal sections (*top*) and its representation by serial cylindrical compartments (*bottom*). Scattered dots illustrate a snapshot of the Monte Carlo simulation test (monitored live in ASTRO; Supplementary Movie 1) in which Brownian particles are injected into the bottom of the 3D structure, and their arrival time at the top is registered, to compare molecular diffusivity (no electric field) and electrodynamic properties (2.5 · 10^3^ V/m electric field in the z-direction, one electron charge *e* = 1.6·10^−19^ C per particle applied) between the two shapes. **g**,The outcome of Monte Carlo tests for the two shapes shown in **e**, for the molecular diffusion flux (top) and ion current (bottom), measured at the top exit of the shapes (as in **e**), upon injection of the Brownian particles into the bottom entry (as in **e**); blue and green, 3D-EM-reconstructed and NEURON-compatible shapes, respectively. See ^53^ regarding details of electrodiffusion simulations.

We therefore developed algorithms and MATLAB-based software (integrated in ASTRO) that enabled us to sample and store small astrocyte processes from our sets of 3D-reconstructed astroglial fragments (Fig. 2a,b; Supplementary ASTRO User Guide p. 12). The 3D EM reconstruction tools used here have been detailed earlier ^10^; ASTRO can use other text-formatted files containing 3D coordinates of the cell membrane ‘mesh’, which is a common outcome of EM 3D reconstruction programs. This procedure aimed at building a database of astroglial nanoscopic structures: each process would comprise a *z*-stack (arbitrarily long) of serial 60 nm sections, with each section represented by reference points scattered on its ‘polygonal’ perimeter (Fig. 2c; Supplementary ASTRO User Guide pp. 12-13).

### Nanoscopic astroglial processes: NEURON-compatible transformation

Our cell modelling approach aimed to take advantage of the NEURON computational environment enabling multiple mechanisms of cellular biophysics. However, NEURON-built cell models employ cylindrical compartments designed to follow the shape of neuronal dendrites or axons. Because astroglial processes have much less regular shapes (Fig. 2a-c), we carried out a separate investigation to establish whether and how cylindrical NEURON compartments can be used to recapitulate geometry and biophysical properties of realistic astrocyte processes.

To attempt the transition from real (3D EM-based) to NEURON-compatible shapes, we transformed individual ‘polygonal’ *z*-stacks of 3D-reconstructed processes into *z*-stacks of cylindrical (disk-shaped) slabs (Fig. 2d). Here, the adjacent polygonal sections with crosssection areas *S*_*i*_ and *S*_*i*+*1*_ which had an intersection area of *T*_*i*_ (Fig. 2d, left and middle), were represented by two ‘main’ cylinder slabs with base areas *S*_*i*_ and *S*_*i*+*1*_ (termed ‘leaves’) plus a ‘transitional’ cylinder slab (termed ‘stalk’) with the base area *T*_*i*_ (Fig. 2d, right). This transformation approximately preserved the ‘diffusion bottleneck’ and the surface-volume relationship of the original shape. Applying this rule systematically transformed 3D-reconstructed nanoscopic processes into the NEURON-compatible, cylinder-based shapes, with the volume and surface areas similar to those of the original shapes (Fig. 2e).

Next, we employed Monte Carlo simulations to systematically compare two key biophysical traits between the original and the NEURON-compatible shapes, the intracellular diffusion transfer rate and the dynamic electrical impedance. These tests involved ‘injecting’ 3000 Brownian particles into one end of the shape and counting their flux rate at the other end, with or without an electric field (Fig. 2f; example in Supplementary Movie 1). The mathematical algorithms involved were detailed, tested and validated against experimental data in our previous studies ^52–54^ In the majority of cases, our tests showed remarkable similarity between the two shapes in their dynamic electrical and diffusion conductance properties (Fig. 2g). In the remaining cases, minor stochastic variation of the cylindrical compartment diameters achieved a similar match.

After we accumulated a representative population of cylindrical compartment radii from the 3D EM measurements (example in Supplementary Fig. 2), we also found that the biophysical match between original and transformed shapes remained when the cylindrical compartments were shuffled randomly with respect to the original shape (example in Supplementary Fig. 3). Thus, the population of nanoscopic astroglial processes could be represented in a NEURON-compatible format based on the statistical properties of 3D-reconstructed cell fragments. The sampling and transformation procedure for nanoscopic processes, as described above, has been integrated in ASTRO for investigators to explore, monitor, and validate (Supplementary ASTRO User Guide, pp. 11-18).

These measurements provided the data set for the astroglial stem tree (Fig. 1d,e) to be populated with nanoscopic processes in accord with their experimentally obtained statistics. However, this stochastically simulated ‘morphogenesis’ had to be further constrained by the experimentally observed tissue-filling properties of astroglia, as described below.

### Tissue volume fraction and astrocyte surface-to-volume ratios

The tissue volume fraction (VF) occupied by astroglia in the hippocampal neuropil has been estimated to range between 5-10% ^8–10, 55^. VF is a key descriptor of astrocyte morphology because astrocytes occupy adjacent tissue domains, with very little overlap ^11, 56^, while their fine processes fill the local volume in a sponge-like manner ^10, 13, 42^. The distribution of astroglial VF in local tissue thus provides an important guide to the astrocyte architecture.

2PE microscopy provides a straightforward tool to monitor VF of live astroglia *in situ* because the fluorescent signal it collects represents emission from within a thin (~1 μm) layer at the focal plane (Supplementary Fig. 4a). Thus, the signal intensity produced by the dye-filled astroglia scales with the local VF occupied by one or more astroglial processes (Supplementary Fig. 4b). At the same time, the 6-10 μm wide astrocyte soma imaged in the same 1 μm focal plane will correspond to 100% VF occupied by the dye-filled cytosol (at least in its brightest part which is least populated with cell organelles; Supplementary Fig. 4c)^10, 14^. Thus the local-to-somatic fluorescence intensity ratio provides direct readout of astroglial VF (Fig. 3a). We therefore first established the typical distribution of VF within individual cell arbours, as a function of the distance from the soma, for CA1 astrocytes (Fig. 3b; blue dots, mean ± SEM for 13 cells). Next, we adjusted the computer-generated nanoscopic morphology to match this VF distribution (Fig. 3b, red dots).

**Figure 3.**
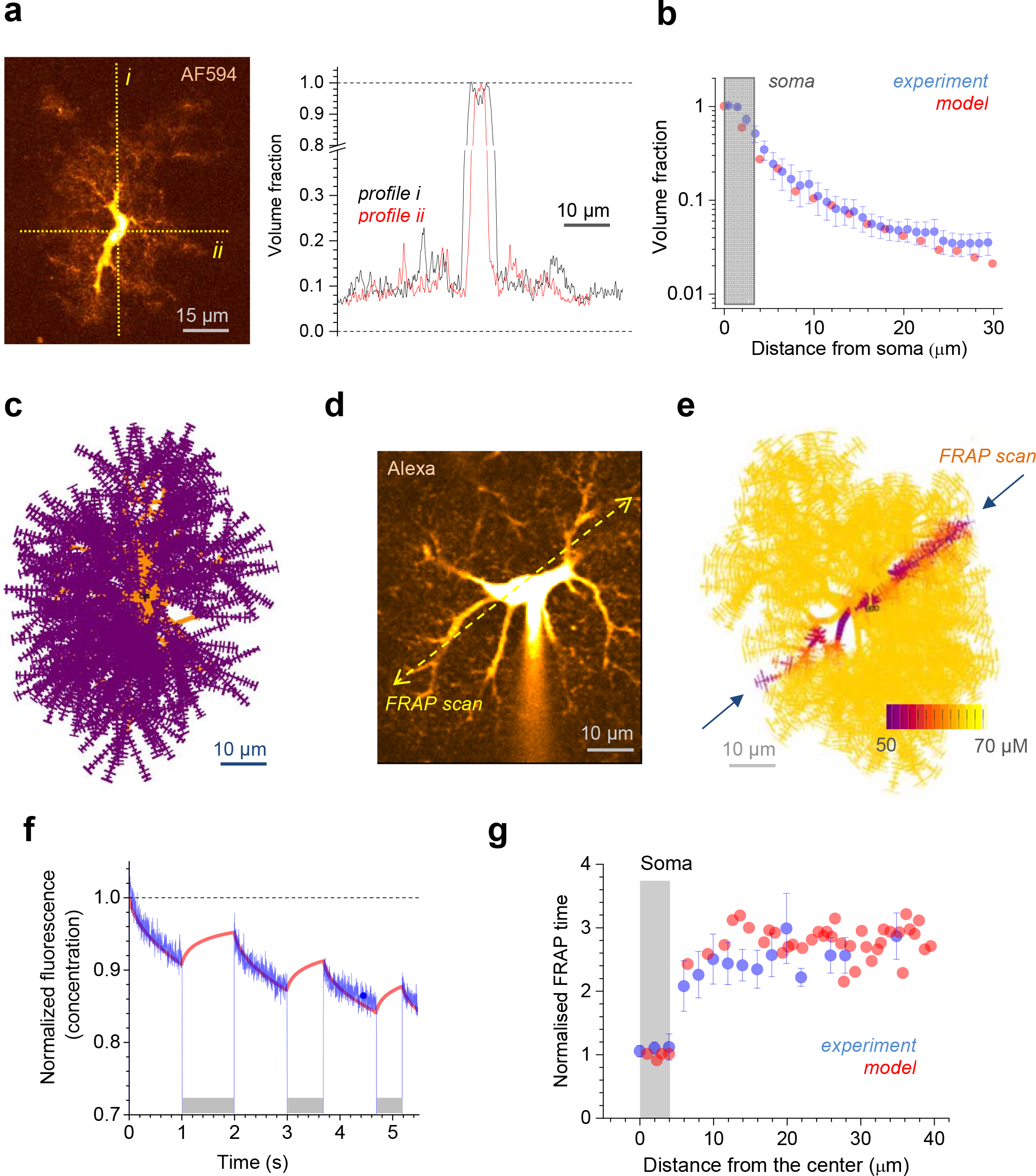
NEURON-based astrocyte model: determining volumetric quantities. **a**, Image panel, a characteristic astrocyte in area CA1 (Alexa Fluor 594, λ_x_^2p^ = 800 nm) seen in a single 2PE optical section (~1 μm thick) crossing the cell soma; dotted lines, sampling fluorescence intensity profiles reporting the astrocyte tissue volume fraction (VF). Graph, VF profiles (fluorescence local/soma ratio) obtained along the dotted lines *i* and *ii* in the image, as indicated. **b**,Matching modelled (red) and experimental (blue; data from n = 13 astrocytes) VF values (ordinate) sampled at different distances from the soma (abscissa). **c**, A complete NEURON-generated astrocyte model (z-projection), with main branches depicted in orange (partly obscured by smaller processes) and nanoscopic protrusions (schematic depiction) in purple. Note that tortuous processes of real-life astroglia are represented by biophysically equivalent ‘straightened’ processes in the model. **d**, An example of astroglia as in a; dotted line, line scan position to measure internal diffusion connectivity (using Alexa Fluor 594 photobleaching); patch pipette is seen. **e**, A snapshot of a photobleaching experiment *in silico* showing the intracellular Alexa concentration dynamics in a modelled astrocyte; arrows, photobleaching line positioning; false colour scale, intracellular Alexa concentration, as indicated (Supplementary Movie 2). **f**, Matching the modelled (red) and the experimental (blue) time course of intracellular Alexa Flour fluorescence during a photobleaching experiment as shown in **d-e**, one-cell example (CA1 area, *stratum radiatum* astrocyte). Grey segments indicate laser shutter-on when fluorescence recovery occurs (red). **g**,Statistical summary of photobleaching experiments (n = 10 astrocytes) and related simulations, as depicted in **d-e**, comparing experimental (blue) and simulated (red) data.

The latter is achieved by systematically adjusting two key structural parameters in the model using ASTRO menus. The first is the average size of nanoscopic processes, determined as the number of leaves per process (up to 100; Supplementary ASTRO User Guide, Fig. 16, p. 19). The second parameter is the ‘seed’ density for nanoscopic processes to be planted on the stem-tree branches, which normally ranges between one to three (SeedNumber variable; Supplementary ASTRO User Guide, p. 20).

Another morphological feature essential to the biophysical identity of cell processes is their surface-to-volume ratio (SVR): this value determines how the transmembrane molecular fluxes are converted into intracellular concentration changes. Careful stereological analyses of hippocampal astroglial processes using 3D quantitative EM arrive consistently at the SVR in the 15-25 μm^−1^ range ^8, 41, 55, 57, 58^. The astrocyte fragments sampled in our experiments (Fig. 2a-c) had SVRs fully consistent with this range, and the modelled nano-shapes (Fig. 2f, Supplementary Fig. 3b) faithfully reproduced this SVR range across the entire simulated cell morphology. The average SVR could be further fine-tuned in ASTRO, by adjusting the distribution of ‘leaves’ and ‘stalks’ in stochastically simulated nanoscopic protrusions (Supplementary ASTRO User Guide, pp. 21-23).

Thus, by systematically generating nanoscopic processes on the ‘typical’ stem tree (Fig. 1e) we arrived at what appears to be a realistic, NEURON-compatible CA1 astrocyte geometry comprising 35000-45000 individual compartments (Fig. 3c; see Supplementary Table 1 for the summary of morphological features pertinent to the present case study of CA1 astroglia). Our final test for the realistic nature of the astrocyte *in silico* was to see if its architecture has intracellular diffusion connectivity similar to that in live astroglia. To address this we carried out a series of experiments and simulations based on fluorescence recovery after photobleaching (FRAP), as outlined below.

### Internal connectivity of astroglia: experiment *versus* model

FRAP protocols have long been used to gauge mobility of fluorescent molecules in live cells: the speed with which fluorescence recovers in a laser-bleached cell region reflects diffusion speed of the fluorescent molecules under study ^59^. Notably, a 2PE FRAP technique applied to water-soluble indicator Alexa Fluor 594 was used to monitor diffusion transport through the ultrathin necks of neuronal dendritic spines ^60^. We sought to apply a similar method to the bulk of nanoscopic astrocyte processes using line-scan FRAP: by bleaching the dye molecules within a thin cylindrical volume across the astrocyte tree, this approach should gauge average diffusion connectivity within the adjacent structures (Fig. 3d; Methods), as shown previously ^61^.

Here, we first confirmed that the intracellular fluorescence showed full recovery well before a 60 s interval (Supplementary Fig. 5a,b), thus indicating no concomitant signal from what could be an immobile dye fraction. Furthermore, 60 s after the first cycle, the second FRAP cycle faithfully reproduced the time course of photobleaching and recovery displayed during the preceding FRAP cycle (Supplementary Fig. 5c). This pointed to the stability of FRAP experiments in the current settings.

We next simulated this experiment in the astrocyte model (Fig. 3e; Supplementary Movie 2) and employed an ‘alternating’ FRAP protocol (with laser shutter repeatedly on-off), to compare simulation outcome with experimental observations. We found that by adjusting one free parameter in the model, the rate of Alexa line-scan photobleaching, it was possible to achieve a good match between experimental and simulated data in individual cells (example in Fig. 3f). The summary (n = 10 recorded cells) also showed a good correspondence between recorded and simulated data for the FRAP rates at variable distances from the soma (Fig. 3g). These observations indicated that the simulated cell morphology was a good representation of the inner connectivity of real-life CA1 astrocytes.

### Passive electrical properties of astrocytes

Specific membrane conductance *G*_m_ is an important determinant of electrical signal propagation in non-excitable cells, such as protoplasmic astroglia. First, we measured input resistance of live astroglia in whole-cell mode: the average value (mean ± SEM: 2.66 ± 0.31 MΩ, n = 15; Fig. 4a) was readily consistent with the previous measurements ^37, 38^. Next, we pulled outside-out patches, estimated the patch membrane area (using the classical voltage-step method ^62^, Methods) and recorded patch resistance: these measurements gave us the average astrocyte *G*_m_ (mean ± SEM: 0.69 ± 0.18 mS/cm^2^; Fig. 4b). Finally, we asked whether this experimental value of *G*_m_ (Fig. 4b) could be reproduced by an astrocyte model constrained by two other independent experimental parameters, (steady-state) input resistance *R*_*i*_ and total membrane area *S*_*mem*_, in accord with Ohm’s law *G*_*m*_ = (*S*_*mem*_*R*_*i*_)^−1^. Because modelled nanoscopic processes are generated stochastically, we have obtained a small representative sample of the astrocyte in question, by repeating the stochastic reconstruction of tissue-filling nanoscopic processes, in accord with the morphological constraints (Fig. 4c). In the sample, the cell volume (resulting from stochastic simulations) varied well within the range of 3000-4800 μm^3^ characteristic of CA1 astroglia ^11, 55^. In each cell in the sample, the free parameter *G*_m_ was adjusted until the cell *R*_*i*_ matched the measured value (Fig. 4a). This test produced the average *G*_m_ value of 0.78 ± 0.04 mS/cm^2^ (Fig. 4c), which was indistinguishable from the experimental *G_m_* value (Fig. 4b) or the earlier empirical measurements ^18^. Thus we have confirmed that our model reproduces ‘correct’ experimental values of *G*_*m*_ without artificially imposing it: in other words, stochastically generated nanoscopic processes replicate realistic membrane properties of astrocytes.

**Figure 4.**
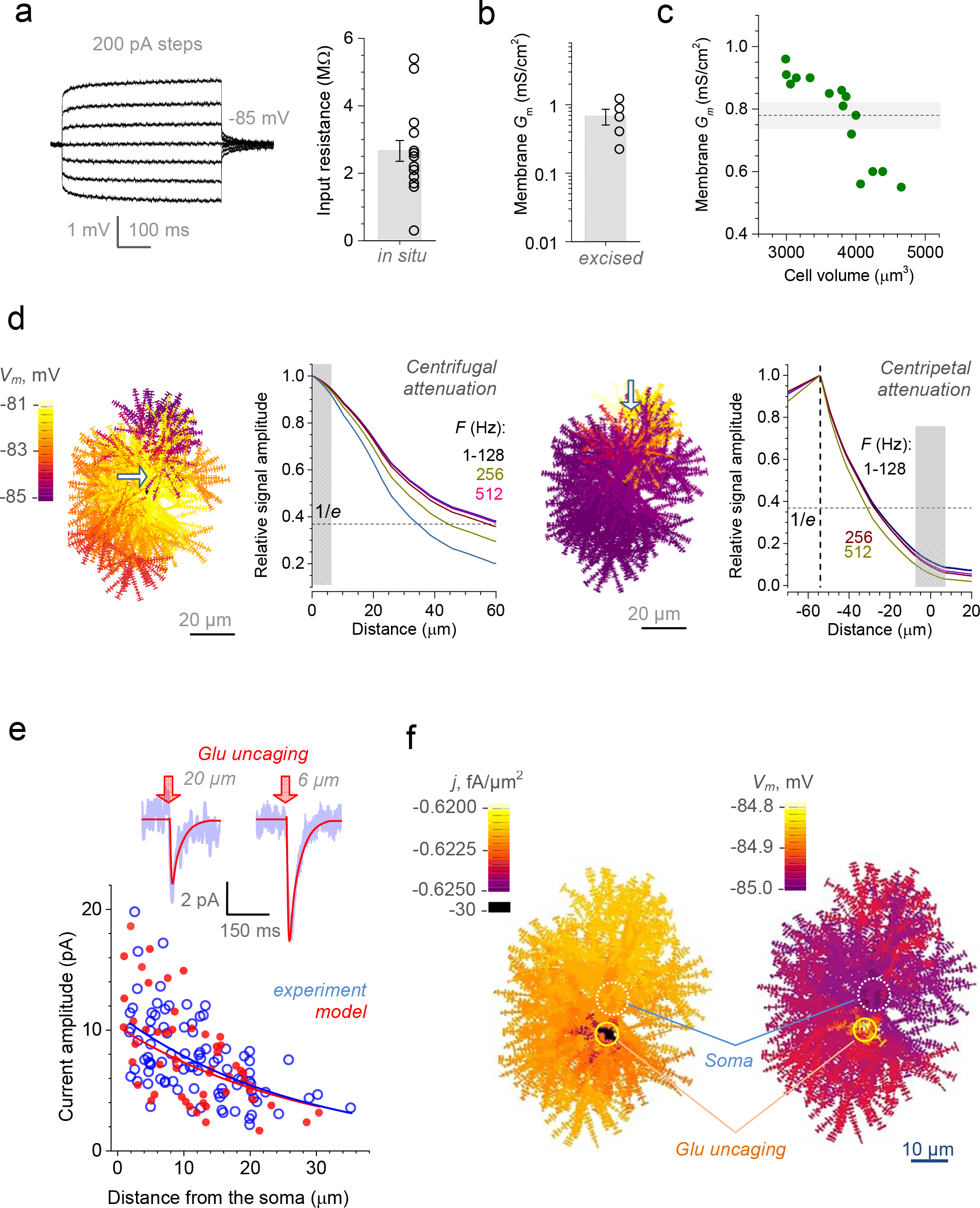
Electrogenic properties of protoplasmic astroglia. **a**, Traces, a characteristic current-voltage recording of CA1 astroglia; graph, input resistance (bar, mean ± SEM; dots, individual cell data; n = 15). **b**,Specific membrane conductance *G*_m_ measured in excised whole-cell (outside-out) patches of CA1 astrocytes (bar, mean ± SEM; dots, individual cell data; n = 5). **c**, Dots, *G*_m_ values obtained by stochastically generating astrocyte models within the empirical range of cell volumes (abscissa) and input resistance matching data shown in **a**; dotted line and grey shade, mean ± SEM for the sample shown; note that NEURON-model astrocyte surface area accounts for both sides and bases of individual cylindrical compartments (Methods). **d**, Membrane space constant estimated using a full astrocyte model for centrifugal (left panels) and centripetal (right panels) voltage signal propagation. Cell shape diagrams: *V*_m_ landscape snapshots generated by local application (shown by arrow) of a sine voltage signal (amplitude +5 mV). Graphs: signal amplitude attenuation at various signal frequencies, as indicated, for centrifugal and centripetal cases, as indicated. **e**, Traces, example of whole-cell recordings (blue) in response to spot-uncaging of glutamate (λ_u_^2p^ = 720 nm, 20 ms duration), at two distances from the astrocyte soma, as indicated; red lines, simulated whole-cell current in the corresponding model arrangement (~5 μm -wide glutamate application; GLT-1 kinetics ^67^; GLT-1 surface density 10^4^ μm^−2^ as estimated ^68^). Plot, a summary of glutamate uncaging experiments (blue open dots, n = 8 cells / 90 uncaging spots) and uncaging tests simulated in the model (red solid dots, n = 39). **f**,Model snapshot 5 ms post glutamate spot-uncaging depicting the cell membrane current (*j*, left) voltage (*V*_m_; right) landscape (example in Supplementary Movie 3); false colour scale.

To understand how far a membrane voltage/current signal propagates in electrically leaky astroglial membranes, we simulated a simple voltage-clamp experiment. This showed that large currents induce very modest astrocyte membrane depolarisation, which propagates with an electrical 3D-space constant of 30-60 μm depending on the signal frequency (Fig. 4d, left). Intriguingly, a membrane voltage signal generated at the periphery is attenuated with a smaller space constant of 25-30 μm (Fig. 4d, right). This asymmetry (reported previously in nerve cells ^63^) arises due to the location-dependent distribution of membrane conductance near the signal cite implying that similar signals arising at different astrocyte loci could have different membrane effects.

Because astrocyte membranes are enriched in potassium channels, in particular K_ir_4.1 type ^17–19^, it would seem important to assess their contribution to the voltage signal propagation. However, the overall unit conductance of K_ir_4.1 in astrocytes is either compatible with or lower than the membrane current leak due to the other conductance contributors (including ion channels, exchangers, gap junctions and semi-channels) ^16, 64^. This suggests that K_ir_4.1 per se has little effect on the passive voltage spread profile, even though these channels play a major role in maintaining the astrocyte resting membrane potential throughout the astrocyte ^65^. For the sake of simplicity, in our baseline test model the multiple membrane mechanisms that hold astroglial resting membrane potential near −83…−85 mV were represented by a non-specific channel current. In these conditions, when we added evenly distributed K_ir_4.1, the effect on voltage propagation was significant only when the overall K_ir_4.1 conductance exceeded substantially its expected physiological range (Supplementary Fig. 6a). In physiological circumstances, this scenario could be affected by changes in extracellular K^+^ concentration ([K^+^]_out_), and by the poorly understood contributing role of other ion channels and exchangers in the astrocyte membrane (see sections below).

### Membrane voltage-current landscape generated by local glutamate uptake

We next asked how the current generated by glial glutamate transporters GLT-1 in response to local glutamate release (a common physiological scenario) alters membrane potential across the cell. First, we recorded from a CA1 astrocyte (acute hippocampal slices) the current response to two-photon spot-uncaging of glutamate (20 ms) at variable distances from the soma (Fig. 4e, traces). To mimic this experiment in the model, we implemented the GLT-1 transporter kinetics ^66, 67^ and ‘scattered’ the transporters uniformly on the cell surface at an effective density of ~10^4^ 2 68 μm^−2^^68^. In these simulations, glutamate was applied within small spherical areas (radius ~3 μm, duration ~20 ms) at quasi-random distances from the soma *in silico*. An excellent match between the modelled somatic current and whole-cell recording data could be obtained (Fig. 4e, plot) by adjusting one free model parameter, the amount / peak concentration of released glutamate. The model unveiled the dynamic landscape of astrocyte membrane voltage, under somatic voltage clamp (to mimic experimental conditions), which varied only within ~0.2 mV across the entire cell (Fig. 4f; Supplementary Movie 3).

To reiterate on the electric properties of astroglia, these data show quantitatively how a local current hotspot in the electrically leaky astroglial membrane stays highly localised, with little effect on the voltage landscape (Fig. 4f). Because of the low baseline membrane resistance, this scenario remained only weakly affected by adding the voltage-dependent K_ir_4.1 conductance ^16, 64^ unless the latter exceeded its physiological range (Supplementary Fig. 6b-d). Again, this test dissects the role of K_ir_4.1 in otherwise unchanged cell conditions (e.g., resting membrane potential stays unchanged as we artificially add K_ir_4.1) whereas removal of K_ir_4.1 in real cells could lead to complex physiological effects ^65, 69^. In any case, these tests suggest that local glutamate transporter currents on their own cannot significantly depolarise astrocyte membrane away from the region of active transport.

### Potassium uptake and redistribution inside astroglia

Rapid uptake of the neural activity-induced excess of extracellular K^+^ and its intracellular redistribution are essential functions of brain astroglia ^18–20, 46, 70^. Several elegant biophysical models have shed light on the kinetics and the tissue-wide spread of this process ^16, 46, 71^. Modelling with ASTRO should help to understand the detailed spatiotemporal dynamics of K^+^ redistribution inside individual astroglial compartments, on the microscopic scale.

As a proof of concept, we simulated a physiological scenario in which extracellular [K^+^] ([K^+^]_out_) was elevated, from resting 3 mM to 10 mM (level often associated intense local excitatory activity ^72^), inside a 20 μm wide spherical region of the astrocyte-containing tissue, for 2 seconds (Fig. 5a). The cell model was populated with K_ir_4.1 channels with the kinetics described earlier ^16^, and unit conductance of 0.1 mS/cm^2^. Upon [K^+^]_out_ elevation, these channels were activated homogeneously inside the 20 μm area (peak current 0.01 mA/cm^2^; not shown). This prompted local K^+^ entry, leading to a local increase in [K^+^]_in_, which dissipated inside the cell over several seconds after [K^+^]_out_ returned to 3 mM (Fig. 4b). During the period of elevated [K^+^]_out_, the membrane potential landscapes reflected very slight depolarisation generated by K_ir_4.1 (Fig. 5c), not dissimilar to the slight depolarisation arising from local glutamate uptake (Fig. 4f). These data provided a quantitative illustration of extracellular K^+^ buffering and its intracellular redistribution, suggesting that the latter is likely to be controlled by local K^+^ efflux, in particular through K_ir_4.1. Again, although K_ir_4.1 appear to represent a major, potential-forming membrane astrocyte conductance ^16, 64, 65^, and further experimental studies are required to understand whether the effect of other membrane mechanisms has any significant contribution to K^+^ buffering.

**Figure 5.**
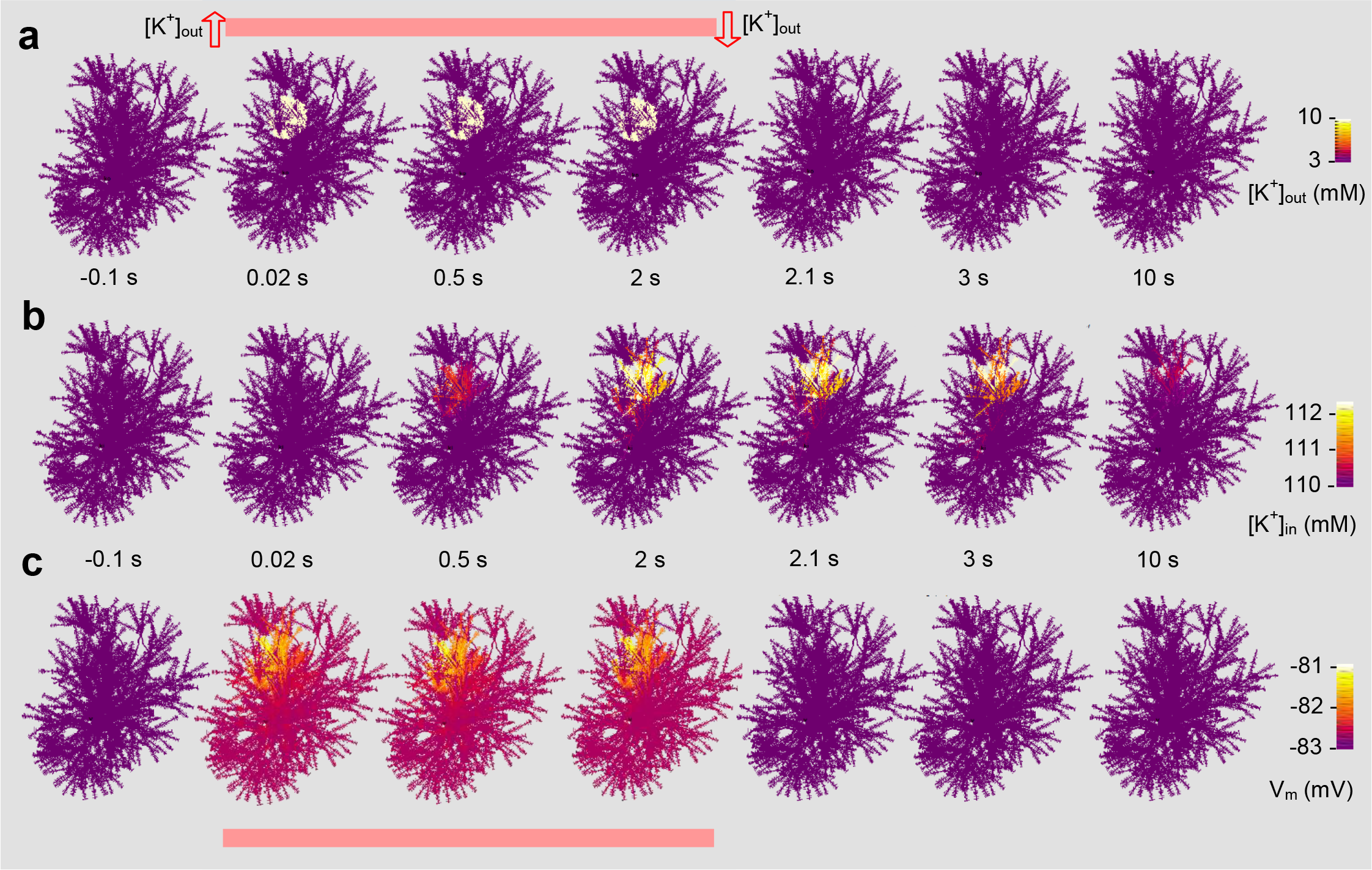
Simulated dynamics of intracellular potassium and membrane voltage triggered by extracellular potassium rise. **a**, Cell shape diagrams, time series snapshots of the cell shape (3D-reconstruction reconstruction shown in Fig. 1a-d) illustrating a spherical 20 μm wide area within which extracellular [K^+^]_out_ was elevated form baseline 3 mM to 10 mM, for 2 seconds (onset at *t* = 0), as indicated; K_ir_4.1 channels were evenly distributed with unit conductance of 0.1 mS/cm^2^ generating peak current density (in the region with [K^+^]_out_ = 10 mM) of 0.01 mA/cm. The K_ir_4.1 kinetics were incorporated in NEURON, in accord with ^16^, as: 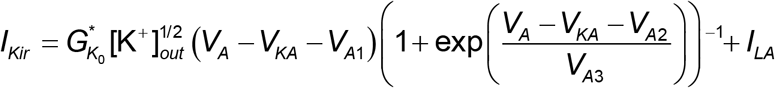 where 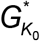 is the effective conductance factor, *V*_ka_ is the Nernst astrocyte K_+_ potential, *V*_*A*_ astrocyte membrane potential, *K*_*0*_ is [K^+^]_out_, *V*_*AI*_ an equilibrium parameter (sets *I*_*kir*_ to 0 at −80 mV), *V*_*A2*_ and *V*_*A3*_ are constants calibrated by the *I-V* curve, and *I*_*LA*_ residual leak current. **b**, Cell shape diagrams, snapshots illustrating the spatiotemporal dynamics of internal [K^+^]_in_ in the test shown in **a**; false colour scale, as indicated. **c**, Snapshots illustrating the spatiotemporal dynamics of the membrane voltage in the test shown in **a**; false colour scale, as indicated.

To further dissect the contributing role of active K^+^ extrusion mechanisms, such as pumps and ion exchangers, we carried out similar tests (K^+^ within a ~15 μm spherical area, over 200 ms at a current density of 1 mA/cm^2^), but with the active removal of intracellular K^+^ by a first-order pump (this may also reflect gap-junction escape) and no contribution from K_ir_4.1 channels (Supplementary Fig. 7). Again, the model captured the dynamic spatial landscape of [K^+^]_in_, with and without any extrusion mechanisms, confirming that local K^+^ efflux could efficiently limit the spatial spread of [K^+^]_in_ elevations (Supplementary Fig. 7).

### Gap junctions and hemichannels

Gap junctions comprising of adjoined connexin proteins connect neighbouring astroglia: this enables electrical current leak and the diffusional flow of molecules across the tissue-wide astroglial syncytium ^40, 73^. In addition, astroglial membranes are enriched in connexin hemichannels permitting transfer of large molecules between the cytoplasm and the extracellular medium ^74^. To understand their contribution to the membrane conductance in our tests, we blocked them with carbenoxolon (CBX, bath applied 50 μm). This increased cell input resistance by ~30% (Supplementary Fig. 8), consistent with the previous measurements in CA1 astrocytes ^75^.

To enable theoretical probing of these features, we have incorporated basic gap junction options in the ASTRO menu, both as electric conductance and as a diffusion channel (Methods; Supplementary ASTRO User Guide, pp. 18, 26-28). For the sake of clarity, however, simulation examples in the present study considered gap junctions and hemichannels as a constituent contributor to membrane conductance. Dissecting their precise functional roles will require further dedicated experimental studies aiming to quantify their biophysical properties as well as their 3D distribution across the astrocyte tree ^76^.

### Astroglial calcium waves: probing the impact of calcium buffering

Astrocytes have long been known to generate non-dissipative Ca^2+^ waves communicating physiological signals within and among cells ^1, 2^. The underlying cellular mechanisms are thought to engage a variety of Ca^2+^ stores, channels, and pumps involving endoplasmic reticulum and mitochondria (reviewed in ^5, 15, 77^). The key molecular machinery here appears to engage Ca^2+^-dependent Ca^2+^ release from Ca^2+^ stores controlled by inositol 1,4,5-trisphosphate (IP_3_) receptors and possibly ryanodine receptors, involving a highly non-linear, sometimes bell-shaped, dependence between their activity and local Ca^2+^ concentration ^78–81^. More recently, significant Ca^2+^ transients have been detected in astroglia lacking IP_3_ receptors ^82–84^, although cell-engulfing, ‘global’ waves seem to involve IP_3_ action (reviewed in ^4^). In all such cases, a critical parameter that controls intracellular Ca^2+^ signal propagation is the Ca^2+^ buffering capacity of the cytosol ^85^, the feature explored in detail in nerve cells ^86^.

The biophysical mechanisms of Ca^2+^ oscillations and Ca^2+^ diffusion and buffering are enabled in the NEURON environment by default, based on the kinetics of IP_3_ signalling which has been well explored (Methods). In addition, ASTRO package incorporates two alternative IP_3_-dependent Ca^2+^ mechanisms described earlier in theoretical studies of astroglial Ca^2+^ signalling ^26, 28, 87^. In the default configuration, four key parameters control intracellular Ca^2+^ wave dynamics: resting (or average) IP_3_ concentration C_IP3_, resting Ca^2+^ level [Ca^2+^]_0_, concentration [*B*] of endogenous Ca^2+^ buffer, and buffer affinity (dissociation constant) *K*_*D*_. Most studies constrain C_IP3_ within 0.5-3 μm ^88–90^. As for [Ca^2+^]_0_, we were able to measure it directly using time-resolved fluorescence microscopy pointing to the range of 50-100 nM ^91^. In contrast, [*B*] and *K*_*D*_ of endogenous Ca^2+^ buffers can vary widely across cell types and thus remain unknown yet essential determinants of intracellular Ca^2+^ dynamics in astroglia.

Similar to the biophysical models of other cell types ^88–90^, we found that Ca^2+^ waves in the modelled astroglia could be generated over a wide (physiologically plausible) range of the aforementioned parameters. To dissect and illustrate the impact of Ca^2+^ buffering on Ca^2+^ wave propagation *per se*, we compared the wave dynamics in modelled cells with and without a relatively small amount (10 μm) of mobile Ca^2+^ buffer added (Fig. 6a; see legend for parameter detail; Supplementary Movie 4). It appeared that adding the buffer reduced the speed and amplitude of [Ca^2+^] waves significantly (Fig. 6a). Intriguingly, this also implies that increased Ca^2+^ buffering in these conditions leads to longer periods of elevated intracellular [Ca^2+^]. Again, an accurate interpretation of such results requires a dedicated experimental study.

**Figure 6.**
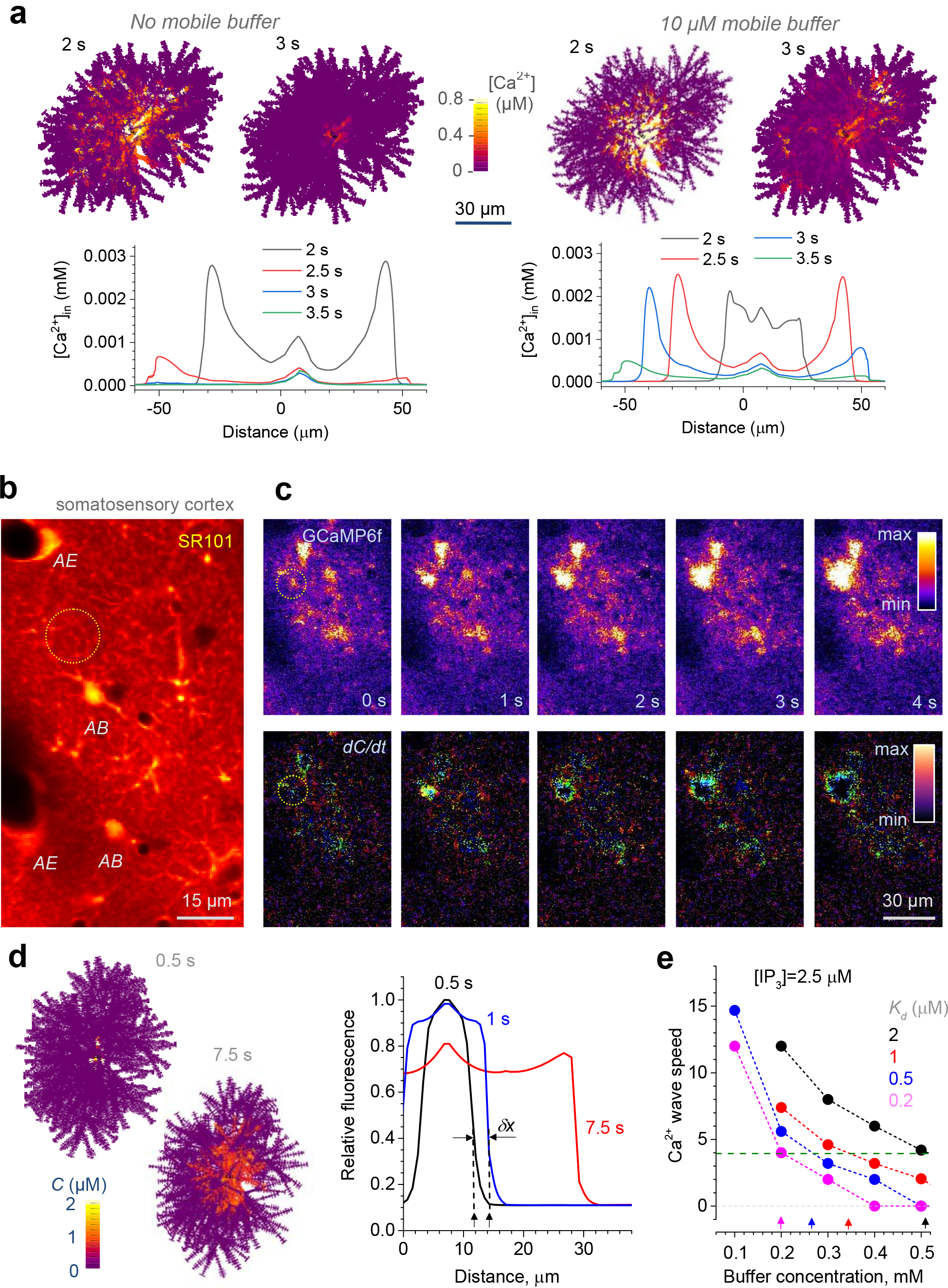
Ca^2+^ waves and Ca^2+^ buffering capacity of astrocytes. **a**, Testing the basic effects of mobile Ca^2+^ buffers on Ca^2+^ wave generation in astroglia; cell diagrams, snapshots of [Ca^2+^] landscape at time points after wave generation, with and without Ca^2+^ buffer, as indicated (note that some branches could obstruct full 3D view); graphs, evolution of the corresponding [Ca^2+^] profiles (zero Distance, soma centre), as indicated. Relevant model parameters: Ca^2+^ diffusion coefficient, 0.3 μm^2^/ms; immobile/endogenous Ca^2+^ buffer concentration, 200 μm (K_f_ = 1000 mM^−1^ ms^−1^; K_D_ = 20 ms^−1^); mobile Ca^2+^ buffer concentration, 10 μm (K_f_ = 600 mM^−1^ m^−1^; K_D_ = 0.5 ms^−1^; diffusion coefficient 0.05 μm^2^/ms); Ca^2+^ pump activation threshold, 50 nM; Ca^2+^ pump flux density, 20 μm/ms; basal IP_3_ concentration, 0.8 μm; IP_3_ concentration upon release, 5 μm; IP_3_ release onset time, 1 s (further detail in Supplementary ASTRO User Guide, Supplementary Movie 4). **b**, 2PE image of the rat somatosensory cortex *in vivo* (~100 μm deep); a single optical section shown with bulk-loaded sulforhodamine 101 (SR101, red channel) to label astroglial structures, as detailed earlier ^91^; *AB*, examples of the astrocyte cell body; *AE*, examples of astrocyte endfoot processes surrounding blood vessels; see Supplementary Movies 1 and 2. **c**, Region of interest (as in b) shown in the [Ca^2+^]-sensitive GGaMP6f (green) channel. *Top*, snapshot sequence (Supplementary Movie 4) depicting generation and spread of an intracellular Ca^2+^ wave (dotted circle); *bottom*, same sequence shown as the time derivative of fluorescence intensity (over 50 ms interval), to highlight the Ca^2+^ wave front; false colour scale. See Supplementary Movie 5 for awake-animal recording example. **d**, Cell shape diagrams, snapshots of the simulated Ca^2+^ wave spreading with the speed that matches experimental observations; false colour scale (C, concentration). Plot, the average intracellular [Ca^2+^] profile depicting the corresponding (centrifugal) Ca^2+^ wave propagation; *δx* illustrates the wave front speed measurement (distance travelled over 0.5 s). **e**, Summary of simulations depicted in c, to determine the cytosol Ca^2+^ buffering capacity (combination of the buffer affinity *K*_*d*_ and its concentration) that would explain the observed Ca^2+^ wave speed; [IP_3_], assumed intracellular concentration of IP_3_ ^88–90^; dotted line, average experimental speed of centrifugal Ca^2+^ waves in astroglia *in vivo*, determined as in (**a-b**) (n = 54 events in approximately 20 individual cells).

### Astroglial calcium waves: assessing cell calcium-buffering capacity *in vivo*

Because there is an ongoing debate on whether astroglial Ca^2+^ waves seen in acute slices or in culture are fully physiological ^92^, we sought to document such waves in live animals, taking advantage of our earlier protocols of astroglial Ca^2+^ monitoring *in vivo* ^91^. However, imaging intracellular Ca^2+^ waves in the hippocampus *in vivo* involves mechanically invasive methods the impact of which on astroglial function is not fully understood. We therefore imaged somatosensory cortex astroglia (accessible with intact brain surface), which appear to show some remarkable similarities in their major morphological features and territorial volumes, across ages, with hippocampal astrocytes ^93^.

We asked therefore whether [*B*] and *K*_*D*_ for cortical astroglia could be assessed by matching the Ca^2+^ wave dynamics documented *in situ* with the modelled outcome. To reduce the concomitants of *ex vivo* preparations and to take advantage of our earlier measurements of astroglial Ca^2+^ *in vivo* ^91^, we recorded spontaneous activity of astrocytes in the rat somatosensory cortex using a virus-transfected Ca^2+^ indicator expressed under a GFAP promoter. To minimise the spatiotemporal filtering effects of free-diffusing Ca^2+^ indicators we used plasma-membrane tethered GCaMP6f, with a ~50 ms Ca^2+^-sensitive fluorescence response time ^94^. The animals were anaesthetized, to limit bursts of sensory input-evoked regional Ca^2+^ rises that could be mistaken for self-propagating Ca^2+^ waves ^95^. Gross morphology of astrocytes was monitored in a red channel using bolus-loading of sulforhodamine 101 (SR101) via a pressurized patch-pipette (Fig. 6b), as described previously ^91^.

In baseline conditions, we could readily detect spontaneous Ca^2+^ waves that appear to engulf entire individual cells (spreading over 10-20 μm in the XY plane with an average frequency of ~5.4 waves a minute over a 160 × 160 μm ROI; Fig. 6c, top panels; Supplementary Movie 5), in good correspondence with previous observations ^96^. The time first derivative of the fluorescence transients could reveal the spreading wave fronts of these signals (Fig. 6c, bottom panels). Thus, only the waves spreading in a clear centrifugal manner (as in Fig. 6c) were selected, to ensure that they represent single-cell regenerative events rather than signals evoked by synchronous external influences.

Our measurements indicated that such waves propagated with an average radial speed of 3.94 ± 0.16 μm/s (mean ± SEM, n = 54 events). Intriguingly, this speed appears significantly lower than that of the stimulus-induced astroglial Ca^2+^ waves in brain slices or in culture (15-25 μm/s, reviewed in ^97^). The relatively fast speed of Ca^2+^ waves in slices or in culture could be because the exogenous stimulus, such as agonist bath or puff application, could prompt a synchronous receptor response over the entire cell expanse. Similarly, in the awake animals, we could detect local waves resembling those under anaesthesia, albeit at a lower frequency (and apparently lower magnitude), in between large synchronous Ca^2+^ elevations engulfing entire regions (Supplementary Movie 6). Again, the possible overlap of such prevalent synchronous events and ‘genuine’ one-cell waves made the anesthetised animals a preferred choice to assess Ca^2+^ buffering properties of astroglia.

In our tests, intracellular Ca^2+^ waves reported by the GCaMP6f fluorescence could be readily reproduced in an astrocyte model in response to an instantaneous Ca^2+^ rise (5 μm for 0.1 ms) near the soma (Fig. 6d). Although we could observe simulated waves over a wide range of buffering parameters, the experimentally documented propagation speed corresponded to only certain values of [*B*] and *K*_*D*_ (Fig. 6e): for the plausible *K*_*D*_ range of 0.2, 0.5, 1 and 2 μm the corresponding [*B*] were 200, 260, 340, and 510 μm, respectively. The empirical relationship between these [*B*] and *K*_*D*_ values was almost perfectly linear (in μm): [*B*] = 170(1 + *K*_*D*_). This simple formula captures the buffering properties of cortical astrocytes, also suggesting the theoretical lower-limit Ca^2+^ buffer concentration of ~170 μm (ignoring the residual buffering effect of membrane-bound GCaMP6f).

### Decoding astroglial Ca^2+^ signals: from recorded fluorescence to the underlying Ca^2+^ dynamics

Historically, the relatively slow, global Ca^2+^ elevations (as in Fig. 6c) have been the most prominent indicator of astroglial activity. Recent advances in high-sensitivity Ca^2+^ imaging have revealed faster and more local Ca^2+^ signals which are prevalent in small processes and probably engage compartmentalised and diverse cell functions (reviewed in ^4, 15, 98, 99)^. In most cases, such observations rely on high-affinity Ca^2+^ indicators, thus providing only a crude reference to the underlying Ca^2+^ dynamics ^14^. To translate recorded fluorescence into the intracellular Ca^2+^ dynamics, one has to model *in silico* Ca^2+^ entry, diffusion, and the buffering by endogenous proteins and Ca^2+^ indicators, as shown in numerous studies of nerve and muscle cells (reviewed in ^18, 100–102^).

We therefore asked to what degree Ca^2+^ imaging data commonly recorded in astroglia *in situ* shed light on the underlying intracellular Ca^2+^ dynamics. First, we imaged CA1 astrocytes loaded whole-cell with Fluo-4 in acute hippocampal slices: these cells showed robust spontaneous Ca^2+^ activity consistent with previously reported data ^12, 38, 103^ (Fig. 7a,b; Supplementary Movie 7).

**Figure 7.**
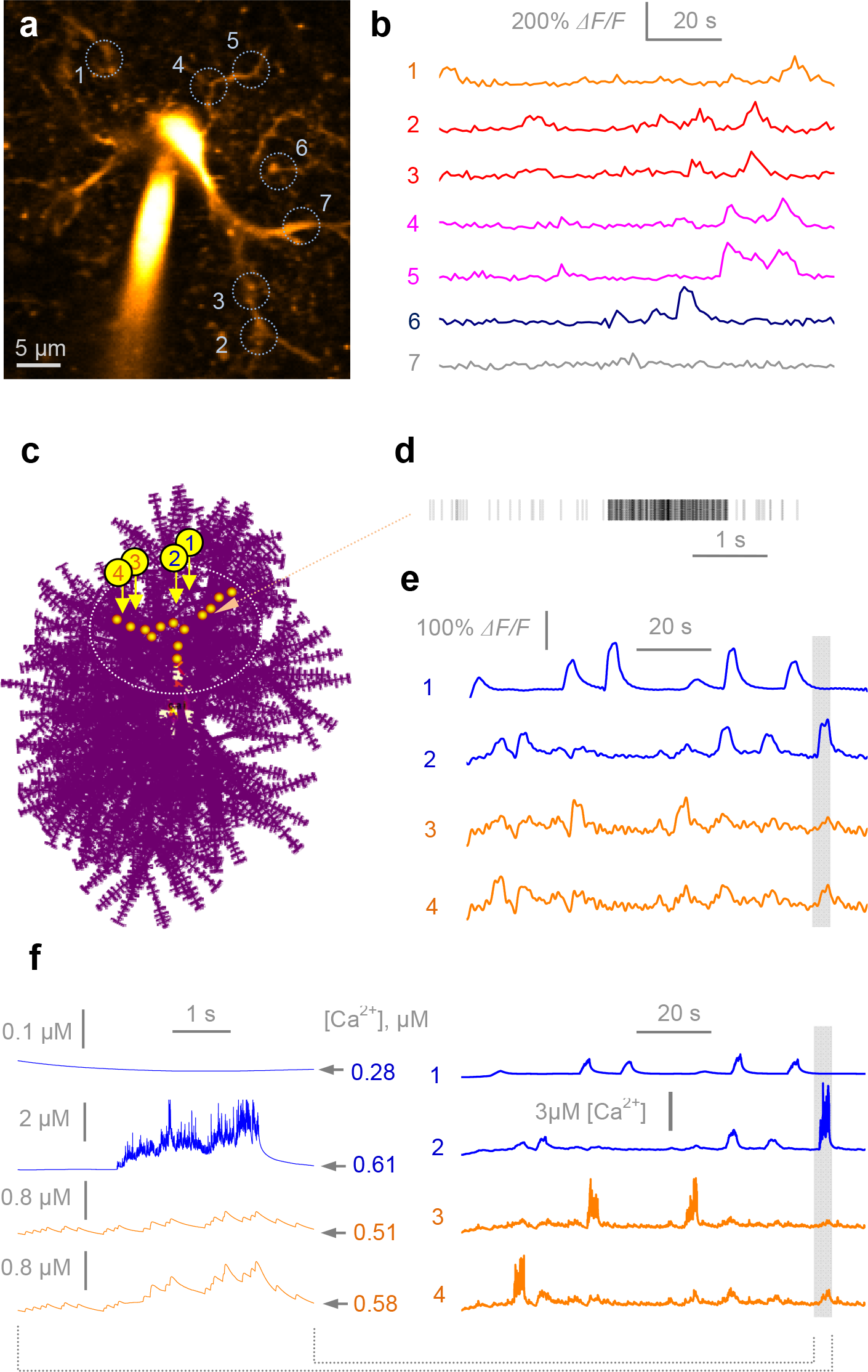
Deciphering intracellular free [Ca^2+^] dynamics from fluorescence Ca^2+^ imaging in astroglia *in situ*. **a**, Example, astrocyte (CA1 area, acute hippocampal slice; Fluo 4 channel, λ_x_ ^2P^ = 800 nm) held in whole-cell, with regions of interest for Ca^2+^ monitoring (circles, ROIs 1-7; Supplementary Movie 7). **b**,Time course of Ca^2+^ sensitive fluorescence (Fluo-4 channel) recorded in ROIs 1-7 as in **a**, over 100 s; same colours correspond to ROIs on the same branch (ROIs 2-3 and 4-5). **c**, An astrocyte model with localised Ca^2+^-puff sources (orange dots) and four recording points (arrows, 1-4); dotted oval, region for analyses: cell area outside has a negligible effect of the Ca^2+^ sources as shown (Supplementary Movie 8). **d**, Example of channel-like local Ca^2+^ entry activity generated by a single localized Ca^2+^ source, in accord with the known biophysical properties of cellular Ca^2+^ sparks and hotspots ^30, 104, 109^. **e**, Time course of simulated Fluo-4 fluorescence (150 μm ‘added’) in ROIs 1-4 shown in **c**: it has statistical properties similar to those recorded *in situ* (**b**); shaded area, time window for higher temporal resolution (see **f**); same line colours correspond to ROIs on the same cell branch. **f**, *Right* simulated intracellular [Ca^2+^] dynamics underlying Fluo-4 fluorescence shown in **e**; *left*, trace fragments on the expanded time scale (shaded area in **e**), as indicated; the fragments correspond to the period of relatively high [Ca^2+^].

While the precise origin of astrocyte Ca^2+^ hotspots is still poorly understood, there has been a large body of work exploring the basic machinery of Ca^2+^ oscillations for other cell types ^80, 81, 104^. The key features emerging from these studies are the occurrence of detectable microscopic Ca^2+^ domains (0.5-5 μm apart ^80, 105^) represented by clusters of stochastically activated Ca^2+^ channels (such as IP_3_ or ryanodine receptors), with the agonist-dependent mean opening time of 2-20 ms, peak amplitude of ~5 pA, and an inter-opening interval varying between zero (overlap, multi-channel opening) and ~50 ms ^106–108^. Stochastic activation of this system produces local Ca^2+^ ‘sparks’ (associated with ryanodine receptors clusters) or ‘puffs’ (associated with IP_3_-receptor clusters) ^80^, typically with a puff event frequency of 0.1-2 Hz ^30, 109^, leading, in certain conditions, to formation of global Ca^2+^ activity ^104^. Genetic deletion of astroglial IP_3_ receptors has unveiled other important sources of spontaneous Ca^2+^ activity, such as mitochondria and Ca^2+^ channels ^82–84^. However, the exact kinetic properties of such Ca^2+^ sources remain to be established.

In our example, in keeping with the basic paradigm, we have scattered local clusters of Ca^2+^ channels (IP_3_ receptor type) along several branches of the modelled astrocyte, 1-5 μm apart (Fig. 7c), within a ~20 μm area of interest: Ca^2+^ activity outside the area was of little consequence because of the comprehensive diffusional dissipation in the absence of global Ca^2+^ events. The simulated astrocyte was ‘filled’ with 150 μm freely diffusing Fluo-4 (*k*_*on*_ = 600 mM^−1^ms^−1^, *k*_*off*_ = 21 ms^−1^) and the endogenous buffer in accord with our estimation (Fig. 6d, we used 200 μm, *K*_*D*_ = 0.2 μm; other combinations consistent with Fig. 6d data produced similar results).

Next, we explored simulated channel openings, within and among Ca^2+^ channel clusters, over the plausible range of their characteristic frequencies (see above; illustrated in Fig. 7d). Astrocyte Ca^2+^ fluorescence was represented by the concentration of Ca-bound Fluo-4 and recorded at four arbitrarily selected points in the area of interest (Fig. 7c; Supplementary Movie 8). We found that at the average interval between channel openings of ~3 ms within clusters, and ~7 s among clusters, the simulated fluorescence signals (Fig. 7e) were similar to the experimental recordings (Fig. 7b). The intracellular Ca^2+^ dynamics underlying these signals was readily available in the model (Fig. 7f). Clearly, obtaining accurate data on the intracellular Ca^2+^ dynamics would require systematic experimental probing involving varied Ca^2+^ buffering conditions and a molecular dissection of the Ca^2+^ cascades involved. However, the example shown here indicates that the classical molecular machinery of Ca^2+^ signalling long explored in other cell types ^80, 81, 104–106, 108^ could generate intracellular Ca^2+^ activity consistent with Ca^2+^ imaging data collected in astrocytes. Our further exploration of IP_3_ arrangement within local astrocyte branches revealed a complex relationship between inter-cluster distances, spontaneous Ca^2+^ activity and its fluorescent-indicator readout (see below).

### Variations in cell features: probing an impact on astroglial function

One of the advantages of having a realistic cell model is the possibility to dissect an impact of specific cellular features or physiological environment (which might be inaccessible for experimental manipulation) on cell’s behaviour. This strategy could also reveal whether certain feature combinations make the modelled cell unstable. To illustrate such an approach, we carried out a selection of example tests in which some key functional traits of modelled cells were monitored against changes in model parameters.

First, to see how strongly gross morphology of the astrocyte could influence its biophysics, we compared two different modelled cells, one with the stem tree reconstructed from an *in situ* experiment (Fig. 1a-d), and the ‘typical CA1 astrocyte’, with the tree branches adjusted to match the average features of CA1 astrocytes (Fig. 1e). The two modelled cells featured different stem trees but were populated with nanoscopic processes based on the same volume fraction, size, and shape distributions (as illustrated in Figs 2-3). Simulations revealed only subtle differences between the cells in their passive voltage spread, input resistance, or Ca^2+^ wave generation (Supplementary Fig. 9). In a similar context, simulating astrocyte ‘swelling’ by ~20% (by evenly increasing the width of cell processes throughout) had only moderate consequences (Supplementary Fig. 10). These examples suggest a relatively narrow range of effects arising from morphological variations *per se*, when all other features remain unchanged.

We next asked how the effective intracellular diffusion coefficient - for instance, for molecules dialysed in whole-cell mode - would affect the molecule equilibration time in the astrocyte. Our simulations mimicking whole-cell dialysis through the soma provide a quantitative illustration of how reducing diffusivity from 0.6 to 0.05 μm^2^ /ms (values characterising obstacle-free diffusion of small ions and 2-3kDa molecules, such as Alexa 488-dextran, respectively) slows down dye equilibration (Supplementary Fig. 11). These examples, however, should be further tested against experimental data, mainly because larger molecules tend to undergo significant steric and viscous hindrance inside small cell compartments (hence diffuse disproportionately slower) ^110^. Also, these example data ignore the possible diffusion escape of the smaller molecules through gap junctions: this feature could be added through the ASTRO menu (Supplementary ASTRO User Guide, pp. 27-28) once the gap junction sink parameters have been experimentally constrained.

Finally, we used simulation settings described in Fig. 7 to ask how the clustering of the IP_3_ dependent Ca^2+^-signalling mechanisms affects local Ca^2+^ activity in astrocyte branches. It appears that spreading this signalling mechanism into individual (equally spaced) clusters, with the same total amount of the IP_3_ activity, facilitates triggering of significant de novo Ca^2+^ events, which could feature prominently in fluorescent recordings (Supplementary Fig. 12).

## DISCUSSION

### Realistic astrocyte models: rationale

The present study sought to create a simulation tool ASTRO that would allow scientists to explore and test mechanistic hypotheses pertinent to complex astroglial physiology, on the scale from nanoscopic processes to the entire cell expanse. Biophysical cell models with realistic morphology have been important for a better understanding of neural function yet there have hitherto been no similar tools available to study astroglia. We therefore aimed at filling this knowledge gap.

In particular, we tried to show specific examples where the ASTRO built model could evaluate how fast molecular diffusion exchange occurs within the astrocyte; whether and how local electric events can control voltage-sensitive membrane mechanisms across the astrocyte; how fast ions such as K^+^ could enter the cell through known channels and how fast they could equilibrate inside it; what key cellular parameters control astrocyte Ca^2+^ waves; how one should interpret Ca^2+^ imaging recordings.

### Building astrocyte models with ASTRO

The task of recreating astroglial morphology *in silico* included three steps. The first step was to construct a ‘stem tree’ based on experimental documentation of main astrocyte branches that are readily identifiable in the microscope. This procedure is similar to the commonly used 3D reconstructions of fluorescently labelled nerve cells using *z*-stacks of their optical sections. The second step, which was the key methodological challenge here, was to recreate the complex morphology of numerous nanoscopic astroglial processes that pervade the synaptic neuropil. To address this, we developed algorithms and computational tools with the aim, firstly, to document and quantify such processes using an empirical 3D EM database, and secondly, to transform the recorded shapes into NEURON-compatible, cylinder-compartment-based shapes, with matched biophysical properties. The latter match was to be verified using specific Monte Carlo simulation tests (for diffusion and electrodiffusion) incorporated in the software. The recorded shapes provided all the key statistics characterising the shapes of nanoscopic astroglial processes in the model.

Thus, the third step was to populate the modelled stem tree with the stochastically generated nanoscopic processes, in accord with their morphometric characteristics obtained as outlined above. The essential experimental constraint here was the tissue volume fraction occupied by astroglial processes, which we and others could measure directly using either 2PE microscopy or 3D EM. Because neighbouring astrocytes display a negligible overlap in the tissue volume, this quantity faithfully reflects the space-filling properties of individual astroglia. We therefore stochastically generated individual processes on dendritic branches until their bulk, within the cell expanse, matched the empirically measured astroglial tissue volume fraction. This procedure completed the architecture of the modelled astrocyte: the model could now be explored using NEURON simulation environment which we equipped with several additional functions specific to astroglial exploration.

In our case study, we obtained and incorporated detailed morphological data on hippocampal astroglia in area CA1, recreated the ‘typical’ cell *in silico*, and explored it against experimental observations involving patch-clamp electrophysiology and 2PE imaging *in situ*. This investigation has shed light on some features of astroglial physiology that have not been attainable in experimental measurements. Our simulations predict that membrane local glutamate uptake or K^+^ intake via K_ir_4.1 generate only very small membrane depolarisation across the astrocyte. It appears that transient rises of extracellular [K^+^] lead to relatively small relative changes of intra-astroglial [K^+^], which dissipate relatively quickly, within one cell, helped by efficient local K^+^ efflux through K_ir_4.1 channels. Our tests illustrated that relatively small changes in the Ca^2+^ buffering properties of astroglia might lead to significant changes in the spread of regenerative intracellular Ca^2+^ signals. The modelling also suggested that the classical molecular mechanisms of rapid Ca^2+^ sparks and hotspots described in other cell types could be consistent with the experimental observations of the apparently slow Ca^2+^ activity documented using common Ca^2+^-sensitive fluorescent indicators.

### Exploring astrocyte function and astroglia-synapse exchange

In broad terms, the astrocyte modelling approach presented here serves several general purposes. Firstly, to recapitulate complex astrocyte morphology at multiple scales, thus providing a first structurally realistic representation of these cells, as well as the possibility to explore the effects of morphological changes on cell function. Secondly, to assess whether experimental results obtained in astroglia, their interpretation or the underlying hypotheses are biophysically plausible, by reproducing them in the realistic model. Thirdly, to understand microscopic spatiotemporal profiles of ion currents, molecular fluxes, and other events that cannot be monitored or registered experimentally. Finally, to predict the relationships between specific cellular features (morphology, Ca^2+^ buffering, channel current density, molecular transport, etc.) and physiological phenotype registered experimentally.

ASTRO has direct access to the full library of biophysical mechanisms pertinent to synaptic and non-synaptic receptors enabled by NEURON. Therefore, arbitrary patterns of synaptic influences throughout the astrocyte tree could be simulated. In addition, excitatory synaptic function could be mimicked using the glutamate uncaging menu in ASTRO. Clearly, the investigator is supposed to incorporate a specific, experimentally constrained kinetic mechanism of the synaptically-induced receptor action of interest, be this IP_3_ release, Ca_2+_ entry, K^+^ fluxes, etc. Judging by the numerous examples from neuronal physiology, each such quest will require a separate, dedicated study combining experimental tests with theoretical probing in the model.

Thus, it is important to stress that our main aim here was not to present one fixed astrocyte model. Instead, we sought to create an interactive modelling tool ASTRO that would enable researchers to test biophysical causality of their experimental observations in astroglial types of interest. The examples presented here provide illustrations of how such tasks could be accomplished. In words, ASTRO is not a ‘frozen’ tool: as new investigatory tasks emerge, it is to be upgraded and equipped with further modelling instruments. The current version of ASTRO and its detailed user’s manual are accessible for download and exploration at https://github.com/LeonidSavtchenko/Astro.

## AUTHORS CONTRIBUTIONS

D.A.R., L.P.S. and C.H. conceived the study; L.P.S. developed and implemented the modelling approach and its computing environment; C.H. designed and carried out morphometric studies *ex vivo*; C.H. and L.B. carried out patch-clamp and imaging experiments and analyses *ex vivo*; T.J. carried out some imaging experiments *ex vivo*, J.P.R. carried out imaging experiments *in* vivo; I.K., M.M., and M.G.S. designed and carried our quantitative EM studies; D.A.R. narrated the study, carried out 3D cell reconstructions, analyses and wrote the paper which was subsequently contributed to by all the authors.

## ACKNOWLEDGEMENTS

This study was supported by Wellcome Trust Principal Fellowship (101896), European Research Council Advance Grant (323113-NETSIGNAL), European Commission FP7 ITN Extrabrain (606950 EXTRABRAIN) (D.A.R.), Russian Science Foundation grant 15-14-30000 (Fig. 3AB data; D.A.R.); German Research Foundation (DFG, SFB1089 B03, SPP1757 HE6949/1 and HE6949/3), the European Commission ITN EU-Glia, Human Frontiers Science Program, NRW Rückkehrerprogramm, UCL Excellence Bridging Award (C.H.). The authors thank Sergey Alexin and Volodymyr Hromakov (AMCBridge LLC) for inspirational help with software solutions. The authors declare no conflict of interest.

## METHODS

### Experimental methods: Electrophysiology *ex vivo*

Acute hippocampal transverse slices (350 μm thick) were prepared from P21-28 Sprague-Dawley rats, in full compliance with the national guidelines, the European Communities Council Directive of 24 November 1986 and the European Directive 2010/63/EU on the Protection of Animals used for Scientific Purposes. Slices were prepared in an ice-cold slicing solution containing (in mM): NaCl 60, sucrose 105, NaHCO3 26, KCl 2.5, NaH2PO4 1.25, MgCl2 7, CaCl2 0.5, glucose 11, ascorbic acid 1.3 and sodium pyruvate 3 (osmolarity 300-310 mOsM), stored in the slicing solution at 34 °C for 15 min and transferred for storage in an extracellular solution containing (in mM): NaCl 125, NaHCO3 26, KCl 2.5, NaH2PO4 1.25, MgSO4 1.3, CaCl2 2 and glucose 16 (osmolarity 300-305 mOsm). All solutions were continuously bubbled with 95% O2/5% CO2. Slices were allowed to rest for at least 60 min before recordings started.

Whole-cell patch-clamp recordings of *stratum radiatum* astroglia were performed in a submersion-type recording chamber. Slices were superfused with an extracellular solution containing (in mM): NaCl 125, NaHCO3 26, KCl 2.5, NaH2PO4 1.25, MgSO4 1.3, CaCl2 2 and glucose 16 (osmolarity 300-305 mOsm), continuously bubbled with 95% O2/5% CO2. Whole-cell recordings were obtained with patch pipettes (3-5 MΩ) with an intracellular solution containing (in mM): KCH3O3S 135, HEPES 10, Tris-phosphocreatine 10, MgCl2 4, Na2ATP 4, Na3GTP 0.4 (pH adjusted to 7.2 with KOH, osmolarity 290-295 mOsM). The cell impermeable Ca2+ indicator OGB-1 (200 μm unless indicated otherwise; Invitrogen O6806) was added to the internal solution. CA1 pyramidal cells were held at −70mV. Protoplasmic astrocytes located in the stratum radiatum were identified by their small soma size, low resting potential (< −80 mV) and low input resistance (<10MΩ). Astrocytes were held in voltage clamp at their resting potential or in current clamp.

In some experiments, whole-cell patches were excised by pulling gently the patch pipette attached to the cell body until the patch was completely detached from the processes and its membrane sealed. Estimation of the patch capacitance *c* was carried out using a classical voltage-step method in which (a) a brief voltage step *ΔV* is applied, (b) the area under the transient capacitance current after the end of the voltage step is measured, giving electric charge *Q*, and (c) membrane patch capacitance is estimated as *c* = *Q* / *ΔV* Thus the patch area is evaluated from the ratio *c* / *c*_m_ where specific membrane capacitance *c*_m_ = 1 μF / cm^2^ is a common characteristic of astroglial membranes^18^. The steady-state current response to the voltage step was used to calculate the patch conductance, which was then normalized to the membrane area to obtain G_m_.

### Experimental methods: 2PE imaging, uncaging, and FRAP *ex vivo*

Astrocytes were filled via whole-cell patch clamp with 40-100 μm Alexa Fluor 594 for 15-20 min. We used an Olympus FV1000 imaging system optically linked to a femtosecond pulse Ti-sapphire MaiTai laser (Newport Spectra-physics). Cells were imaged using an Olympus XLPlan N 25x water immersion objective. Fluorescence recordings were obtained in line-scan mode (500Hz, line placed through the astrocyte arbour and across the soma) at λ = 800 nm at an increased laser power of 15-20 mW under the objective to induce substantial bleaching of Alexa Fluor 594. Fluorescence was collected for 750-1000 ms, then a mechanical shutter was placed in front of the laser beam for 1-2 s to allow fluorescence recovery.

We used a combined two-photon uncaging and imaging microscope (Olympus, FV-1000MPE) powered by two Ti:Sapphire pulsed lasers (Chameleon, Coherent, tuned to 720 nm for uncaging and MaiTai, Spectra Physics, tuned to 800 nm for imaging). Cells were imaged using an Olympus XLPlan N 25x water immersion objective. The intensity of the imaging and uncaging laser beams under the objective was set to 5 mW and 12-17 mW, respectively.

To record spontaneous Ca2+ transients in frame scan mode, 200 μm Fluo-4 (Invitrogen) and 100 μm Alexa Fluor 594 (Invitrogen) were added to the intracellular solution. 350-500 μm^2^ fields of view where imaged within the arbour of the patched astrocyte and the fluorescence emitted by Alexa Fluor 594 and Fluo-4 was collected at a rate of 3-5 Hz for 2-3 min. Time-dependent fluorescence transients were expressed as ΔG/R where G corresponds to the background-subtracted Fluo-4 fluorescence and R to the background-subtracted Alexa Fluor 594 fluorescence.

For MNI-glutamate uncaging, astrocytes were loaded with 100 μm Alexa Fluor 594 as a morphological marker. Astrocytes were held in voltage-clamp mode at their resting membrane potential (typically between −80 and −90 mV). The MNI-glutamate (12.5 mM) was either puffed within the tissue from a glass pipette placed above the patched cell, or added to the bath at 2.5 mM. Glutamate was uncaged for 20 ms at different distances from the soma (5-25 μm).

### Experimental methods: 3D reconstruction of live astrocyte stem tree

A *stratum radiatum* astrocyte was held in whole-cell mode, with Alexa Fluor 594 added to the intracellular solution (see above; excitation at λ_x_^2p^ = 800 nm). A *z*-stack of 2PE images was collected using 100 × 100 μm (512 μ 512 pixel) individual frames containing the entire visible astrocyte structure, with a 0.5 μm *z*-step over 61 μm in depth. The image stack was stored (8-bit tiff format), individual images were corrected for the depth-dependent, quasi-exponential fluorescence signal decrease (Fiji Image-Adjust-Bleach Correction, plugin by Kota Miura 2014: 10.5281/zenodo.30769). Fluorescence background was subtracted (Fiji Image-Process), identifiable cell branches (> 0.3-0.5 μm in diameter) were traced semi-automatically in individual 2D optical sections and reconstructed in 3D using Neurite Tracer (Fiji Plugins-Segmentation-Simple Neurite Tracer; by Mark Longhair and Tiago Ferreira, MRC and Janelia Campus; http://imagej.net/Simple_Neurite_Tracer; default segmentation sigma, 0.196). The data sets representing diameters of tubular compartments and their 3D co-ordinates (pairs of end points) were stores in SWC format. The Vaa3D software (Allen Institute, http://www.alleninstitute.org/what-we-do/brain-science/research/products-tools/vaa3d/) was used to convert these data sets into NEURON compatible files providing 3D structure of the astroglia stem-tree (with tubular compartments representing individual cylindrical compartments).

### Experimental methods: Astroglia-targeted expression of GCaMP6f *in vivo*

Animal procedures were conducted in accordance with the European Commission Directive (86/609/ EEC) and the United Kingdom Home Office (Scientific Procedures) Act (1986). Young male C57BL/6 mice (2 - 3 weeks of age) were anaesthetised using isoflurane (5% induction, 1.5 - 2.5% v/v). Subcutaneous analgesic (buprenorphine, 60 μg kg^−1^) was administered and the animal was secured in a stereotaxic frame (David Kopf Instruments, CA, USA) and kept warm on a heating blanket. The scalp was shaved and disinfected using three washes of topical chlorhexidine. Upon loss of pedal withdrawal reflexes, a small midline incision was made to expose the skull. A craniotomy of approximately 1 - 2 mm diameter was performed over the right somatosensory cortical region using a high-speed hand drill (Proxxon, Föhren, Germany). Stereotactic coordinates were +0.1 mm on the anterioposterior axis relative to bregma, and 2 mm lateral to midline. Once exposed, a warmed aCSF variant (cortex buffer, in mM; 125 NaCl, 2.5 KCl, 10 HEPES, 10 glucose, 2 CaCl_2_, 2 MgSO_4_) was applied to the skull and cortical surface throughout the procedure.

AAV5 *GfaABC1D-LckGCaMP6f* (Penn Vector Core, PA, USA) was pressure injected into the somatosensory cortex using a pulled glass micropipette stereotactically guided to a depth of 0.6 mm beneath the pial surface, at a rate of approximately 1 nL sec^−1^. A given injection bolus contained between 0.25 and 0.5 × 10^10^ genomic copies, in a volume not exceeding 500 nL. After injection, pipettes were left in place for 5 minutes before retraction. The scalp was sutured with absorbable 7-0 sutures (Ethicon Endo-Surgery GmbH, Norderstedt, Germany) and the animal was left to recover in a heated chamber. Meloxicam (subcutaneous, 1 mg kg^−1^) was administered once daily for up to two days following surgery. After a 4 - 6 week AAV incubation period, animals were prepared for multiphoton imaging through a cranial window implantation as described below.

### Experimental methods: Two-photon excitation imaging of astroglia *in vivo*

Following viral transduction of LckGCaMP6f as above, male C57BL/6 mice (7-9 weeks of age) were prepared for cranial window implantation and 2PE microscopy. Animals were anesthetized using fentanyl, midazolam and medetomidine (i.p., 0.05, 5 and 0.5 mg kg^−1^, respectively). Adequate anesthesia was ensured by continuously checking for the loss of pedal withdrawal reflexes and anaesthesia was supplemented appropriately throughout the procedure (typically 10-20 % of the original dose per hour). Body temperature was maintained at 37.0 ± 0.5 °C using a feedback rectal thermometer and heating blanket. The animal was secured in a stereotaxic frame and a craniotomy of approximately 2.5 mm diameter was carried out over the right somatosensory cortex, centred 0.2 mm caudal to bregma and approximately 2.5 mm laterally from the midline. Once exposed, the cortical surface was continuously superfused with warmed aCSF (in mM; 125 NaCl, 2.5 KCl, 26 NaHCO_3_, 1.25 Na_2_HPO_4_,18 Glucose, 2 CaCl_2_, 2 MgSO_4_; saturated with 95% O_2_ / 5% CO_2_, pH 7.4). Cortical astrocytes were labeled using multicell bolus loading of sulforhodamine 101 (SR101, 5μm). SR101 (in cortex buffer vehicle) was pressure-injected through a pulled glass micropipette targeted to 2 - 3 injection sites within the transduced region, comprising a total volume of 500nL. The cortical surface was covered with 1% agarose and a glass coverslip was placed on top. Using tissue adhesive (Dermafuse, Vet-Tech Solutions, UK), the coverslip was partially secured and a custom-built headplate fixed to the skull. A single cranial-mounted screw was inserted over the contralateral hemisphere and the entire assembly was then secured using dental cement. During imaging, the headplate was used to secure the animal under the objective on a custom-built stage.

In these experiments, two-photon excitation was carried out using a Newport-Spectraphysics Ti:sapphire MaiTai laser pulsing at 80 MHz, and an Olympus FV1000 with XLPlan N 25x water immersion objective (NA 1.05). Acquisitions were carried out using a wavelength of 920 nm and the mean laser power under the objective was kept at 20 - 35 mW. Cortical astrocytes were readily identified through SR101 labeling and verified for GCaMP6f expression by frame-scanning for calcium transient activity. Recordings were made at a depth between 50 and 250 μm from the cortical surface. XY time series (at 0.5 - 2 Hz with a pixel dwell time of 0.5 - 4 ps and pixel size of 0.248 - 1.59 μm) were taken in identified regions to measure spontaneous calcium activity.

### Experimental methods: Fast fixation and DAB staining of recorded astrocytes

In a subset of experiments we loaded an astrocyte with biocytin, and after the experiment the slices were rapidly fixed (by submersion) with 1.25% glutaraldehyde and 2.5% paraformaldehyde in 0.1 M PB (phosphate buffer, pH 7.4), to be kept overnight, submerged in 10% sucrose in PB for 10 min and then in 20 % sucrose in PB for 30 min. The slices were consequentially freeze-thaw in liquid freon and liquid nitrogen for 3 sec each to gently crack intracellular membranes and embedded in 1% low gelling temperature agarose in PB (Sigma-Aldrich, USA). Embedded slices were sectioned at 50 μm on a vibrating microtome (VT1000; Leica, Milton Keynes, UK). 50 μm sections were incubated in 1% H_2_O_2_ in PB for 20 min to eliminate blood background, washed with 0.1 M TBS (tris buffer saline, pH 7.4) and incubated with ABC solution (VECTASTAIN ABC, Vector laboratories, USA) for 30 min at room temperature. Next section were washed with 0.1M TB (tris buffer, pH 7.4), pre-incubated with DAB (3,3’-Diaminobenzidine tablets - Sigma-Aldrich, USA) solution (10 mg DAB tablet + 40 ml TB) for 30 min at room temperature in dark and finally incubated with DAB+ H2O2 solution (5 μl of 33% H_2_O_2_ + 25 ml of DAB solution) for 10-20 min at room temperature in dark. The DAB stained sections was washed in PB, post-fixed in 2% osmium tetroxide and further processing and embedding protocols were essentially similar to those reported previously (Medvedev et al., 2010). Briefly, the tissue was dehydrated in graded aqueous solutions of ethanol (40 - 100%) followed by 3 times in 100% acetone, embedded into a mixture of 50% epoxy resin (Epon 812 / Araldite M) and 50% acetone for 30 min at room temperature, embedded in pure epoxy resin, and polymerized overnight at 80°C. Sections in blocks were coded and all further analyses were carried out blind as to the experimental status of the tissue.

### Experimental methods: 3D electron microscopy

Serial sections (60-70 nm thick) were cut with a Diatome diamond knife as detailed and illustrated earlier ^111, 112^, and systematically collected using Pioloform-coated slot copper grids (each series consist of up to 100 serial sections). Sections were counterstained with 4% uranyl acetate, followed by lead citrate. Finally sections were imaged in *stratum radiatum* area of CA1 (hippocampus) using AMT XR60 12 megapixel camera in JEOL 1400 electron microscope. Serial sections were aligned as JPEG images using SEM align 1.26b (software available from http://synapses.clm.utexas.edu/). 3D reconstructions of DAB stained astrocyte fragments and the adjoined to stained astrocytes dendritic spines (that host clearly identifiable excitatory synapses) were performed in Trace 1.6b software (http://synapses.clm.utexas.edu/). 3D reconstructions of selected astrocytic segments and dendritic spines were imported to 3D-Studio-Max 8 software for rendering of the reconstructed structures.

### Statistics summary

The present study contained no longitudinal or multifactorial experimental designs. In electrophysiological or imaging experiments the main source of biological variance in were either individual cells or individual preparations (the latter in case of field measurements in acute slices), as indicated. In accord with established practice, in the *ex vivo* tests we routinely used one cell per slice per animal, which thus constituted equivalent statistical units in the context of sampling, unless indicated otherwise. Statistical hypotheses pertinent to mean comparisons were tested using a standard two-tailed t-test, unless the sample showed a significant deviation from Normality, in which case non-parametric tests were used as indicated. The null-hypothesis rejection-level was set at α = 0.05, and the statistical power was monitored to ensure that that the sample size and variance were adequate to detect a mean difference (in two-sample comparisons) of 10-15% or less.

### Astrocyte model: generating ‘invisible’ nanoscopic morphology

Nanoscopic processes of the astrocyte model are generated in a probabilistic manner based on the sample statistics from 3D EM reconstructions (Fig. 2). The total cell surface area *S*_*tot*_ represented by the cylinder-based shape approximations (Fig. 2d,e), consists of the (lateral) surface areas of all cylinder-compartment sides *S*_*lat*_ added to the surface areas of ‘main’ cylinder bases *S*_*M*_ (blue in Fig. 2d, bottom) minus the surface areas of ‘transitional’ cylinder bases *S*_*T*_ (green in Fig. 2d, bottom). In our case study, computations indicated that *S*_*T*_ = 0.20*S*_*M*_ throughout modelling: thus, the formula *S*_*tot*_ = *S*_*lat*_ +0.8*S*_*M*_ was applied. In the generated cell models, the S/V ratios were ranging from ~7 μm^−1^ near the soma to an average of ~22 μm^−1^ in the bulk of the cell arbour, in accord with the empirical observations.

### Astrocyte model: transporter/channel kinetics and diffusion-reaction mechanisms

Models built with ASTRO can incorporate many dozens of NEURON-enabled channel and transporter kinetic mechanisms that have been tested and validated in numerous studies combining experiments and simulations^24^. The formal descriptions of the respective algorithms could be found using an extensive NEURON database at https://senselab.med.yale.edu/modeldb/ which also contains references and links to the original studies and the mathematical formulism involved. Upon ASTRO installation on the host computer, these mechanisms could also be inspected in the respective *.mod files in the ‘neuronsims’ directory or, alternatively, online here https://github.com/LeonidSavtchenko/Astro/tree/master/neuronSims.

Several channel current and diffusion-reaction mechanisms have been written specifically for the present model. The kinetics of glutamate transporter GLT-1 involving glutamate and ion fluxes has been incorporated in accordance with ^113^ (description in the GluTrans.mod file). The K_ir_4.1 potassium current has been incorporated in accordance with ^16^ (Kir4.mod; Fig. 5 legend), intracellular K^+^ diffusion was incorporated as longitudinal diffusion (no radial rings) using a built-in Ca^2+^ diffusion algorithm described in the next section (potassium.mod), the FRAP mechanism incorporated the same algorithm plus a reaction-diffusion step (FRAP.mod), and K^+^ extrusion was modelled as a first-order pump (kpump.mod; Supplementary Fig. xx legend). Gap junction mechanisms were enabled either as a (zero-order) current leak (gap.mod) or as a diffuse escape (gapCa.mod). Throughout these mechanisms, the respective kinetic parameters can be set using the relevant NEURON-enabled ASTRO menus, as described in the User Guide (https://github.com/LeonidSavtchenko/Astro/blob/master/ASTROUserGuide.pdf).

### Astrocyte model: Ca^2+^ homeostasis and diffusion

ASTRO simulation algorithms enabling intracellular Ca^2+^ homeostasis and diffusion (including that among adjacent compartments of unequal size) are detailed in Chapter 9 of the NEURON Book^24^ (also here https://www.neuron.yale.edu/neuron/docs), and can be found in the modified cadifus.mod file in the model installation. In brief, Ca^2+^ diffuses freely whereas buffer-bound Ca^2+^ (which has much lower diffusivity) is considered stationary, for the sake of simplicity. In individual cylindrical cell compartments, radial diffusion occurs through four concentric shells surrounding a cylindrical central core, and longitudinal diffusion is calculated using fluxes between the corresponding concentric compartments adjusted for the cross-section areas. The longitudinal and radial diffusion coefficient for Ca^2+^ was set to 0.3 μm^2^ ms^−1^, the basal level was set to 50 nM, and IP_3_ concentration at 0.

In addition to free diffusion, Ca^2+^ homeostasis mechanisms included the SERCA pump, SERCA channel and SERCA leak, the endogenous (stationary) and exogenous (Ca^2+^ indicator) mobile buffers, and a plasma membrane Ca^2+^ pump with the threshold mechanism (cadifus.mod). The kinetics of buffers can be modified using NEURON-enabled menus. The mechanistic details of Ca^2+^ SERCA pump were as described earlier ^87, 114^. The current model implementation assumes that IP_3_ is distributed uniformly across cell compartments, i.e. that diffusion equilibration of IP_3_ is fast compared to Ca^2+^ concentration transients in space or time.

### Modelling with ASTRO: On-line access and installation

Detailed information on the installation and running of ASTRO can be found in the User’s Manual (Supplementary Material; online download at https://github.com/LeonidSavtchenko/Astro/blob/master/Manual). The current version of ASTRO can also be downloaded directly from https://github.com/LeonidSavtchenko/Astro. The
(regularly updated) User Guide can be downloaded from the same location or found (current version) in the Supplementary Information (User_Guide.doc).

In brief, running ASTRO without full-scale simulations of intracellular Ca^2+^ dynamics requires the host computer to have MATLAB (2012 version or later, https://uk.mathworks.com/products/matlab.html) and NEURON (7.2 or later, https://neuron.yale.edu/neuron/download) installed under Windows 7 or Windows 10.

Simulating full intracellular Ca^2+^ dynamics is highly resource-consuming and normally requires an additional Worker computer / cluster operating under Linux, with preinstalled NEURON (https://neuron.yale.edu/neuron/download/compilelinux) and MPI whereas the Host computer will require MATLAB (2013 version or later), NEURON (7.0 or later), and access to the Internet. In house, the Linux version with the parallel computations including intracellular diffusion simulation was routinely run using a 12-node in-house computer cluster ^115^, taking advantage of the computational optimization routines developed by us earlier for compartmental models and Monte Carlo simulations ^52, 53, 115^.

